# Ratiometric signaling produces robust temporal integration for accurate cellular gradient sensing

**DOI:** 10.1101/2025.04.18.649595

**Authors:** Debraj Ghose, James Nolen, Kaiyun Guan, Timothy C. Elston, Daniel J. Lew

**Affiliations:** Wyss Institute for Biologically Inspired Engineering, Harvard University, Boston, MA, 02215, USA; Department of Mathematics, Duke University, Durham, NC, 27708, USA; Curriculum in Bioinformatics and Computational Biology, University of North Carolina at Chapel Hill, Chapel Hill, NC, 27599, USA; Department of Pharmacology and Computational Medicine Program, University of North Carolina at Chapel Hill, Chapel Hill, NC, 27599, USA; Department of Biology, Massachusetts Institute of Technology, Cambridge, MA, 02139, USA

**Author notes:** These authors contributed equally to this work (Debraj Ghose, James Nolen).

## Abstract

Cells excel at interpreting noisy chemical gradients to guide fertilization, development, and immune responses, but the mechanisms underlying this remarkable ability remain poorly understood. Previous work showed that some G protein signaling pathways can overcome challenges from uneven receptor distribution by using a ratiometric signaling strategy. In this mechanism, G proteins receive information from both bound and unbound receptors, unlike classical signaling where only bound receptors contribute. Here, we show that ratiometric signaling also provides an unexpected ability to suppress noise from low receptor numbers. The benefit stems from each G protein remembering the last receptor state it encountered, so that at any instant, ratiometric G protein collectives reflect time-averaged receptor activity. Unlike classical signaling, this averaging remains unbiased and accurate across the varying ligand concentrations present in a spatial gradient. Using theory and simulations, we demonstrate that this averaging mechanism allows cells to surpass theoretical limits for gradient detection from instantaneous receptor information alone. Our findings reveal how ratiometric biochemical architectures enable robust temporal integration across spatially varying signals, providing cells with enhanced directional accuracy under noisy conditions.

## I. INTRODUCTION

Cells are exceptional at extracting directional information from their chemical environment, often under complex and noisy conditions that demand high accuracy. Some of the earliest direct experimental evidence that cells respond to chemical gradients emerged in the late 19th century, with studies involving bacteria clustering near oxygen-rich zones [1, 2] and plant spermatozoids finding female reproductive organs [3]. Since then, chemotactic or chemotropic responses by cells or subcellular components have been documented in pollen tube guidance [4, 5], protozoa [6–8], fungi [9–14], immune cells [15, 16], animal sperm [17– 19], neurons [20, 21], developmental processes [22], cancer cells [23–25], and many other systems. Together, these observations broadly illustrate the diverse scenarios under which gradient-driven cell guidance occurs.

In all cases, cells detect chemical signals through binding to cell-surface receptors that engage internal signaling pathways [26, 27]. Using these receptors, a cell can in principle estimate the direction of a ligand gradient by measuring the difference in chemical concentration across its width. However, the rapid movement of fast-swimming cells like bacteria or sperm causes them to collide with ligand molecules more often at their front than at their back. This differential collision rate can make the concentration gradients they perceive across their width appear hundreds of times steeper than the actual environmental gradients [28]. Such cells have developed “temporal sensing” strategies that monitor concentration changes over time as they move through space. In contrast, a majority of larger eukaryotic cells, which are slower moving or non-motile, employ a “spatial sensing” strategy that compares concentrations of chemicals on different sides of the cell to infer gradient direction. Once a direction is established, eukaryotic cells can polarize and move or grow toward (or away from) higher ligand concentrations. Even so, these spatial measurements can be limited by gradient measurements with low signal to noise, especially for smaller cells. These limits have been theoretically studied for both absolute ligand concentration measurement [28–30] and gradient measurement [31, 32], grounded in considerations of ligand-binding noise. A well-appreciated strategy to enhance low signal is to use time-averaging of receptor signals, which can significantly improve both concentration measurement and gradient detection. However, molecular mechanisms for averaging receptor signals and improving gradient detection remain poorly understood.

In classical G protein signaling, G proteins are activated by ligand-bound receptors and deactivated at a uniform rate. Here, inspired by the mating response of budding yeast, we examined an alternate strategy: ratiometric signaling, in which G proteins are activated by bound receptors and deactivated by unbound receptors [33, 34]. This architecture means that downstream signaling depends on the ratio of active to inactive receptor levels rather than absolute number of active receptors.

We discovered that ratiometric signaling enables robust time averaging that significantly outperforms classical signaling. This mechanism arises from collectives of downstream G proteins that remember past receptor states. While classical signaling fails to maintain accurate averaging across the varying concentrations present in a spatial gradient, ratiometric signaling automatically adjusts to different concentrations. Furthermore, this mechanism allows cells to surpass theoretical limits for instantaneous gradient sensing, revealing how cells achieve their remarkable ability to interpret noisy directional signals. These results reveal design principles underlying robust spatial gradient sensing and suggest strategies for engineering synthetic systems that can navigate complex chemical environments.

## II. RESULTS

### A. Instantaneous limits for gradient estimation by receptors

To evaluate the potential benefits or pitfalls of a ratiometric sensing strategy, we need a metric that tells us how accurately a theoretical cell can infer the direction of a ligand gradient. Previous studies have compared the ligand concentrations sensed by the front and back halves of a cell, thereby determining whether the difference (signal) exceeds the noise [28, 31]. However, splitting the cell in half presupposes prior knowledge of the axis along which concentration changes occur. Because a real cell generally does not have that information, other work has instead estimated the cell’s “best guess” of the gradient direction by summing the vectors from the cell center to all active receptors [32, 35] (Fig. 1A).

**FIG. 1.**
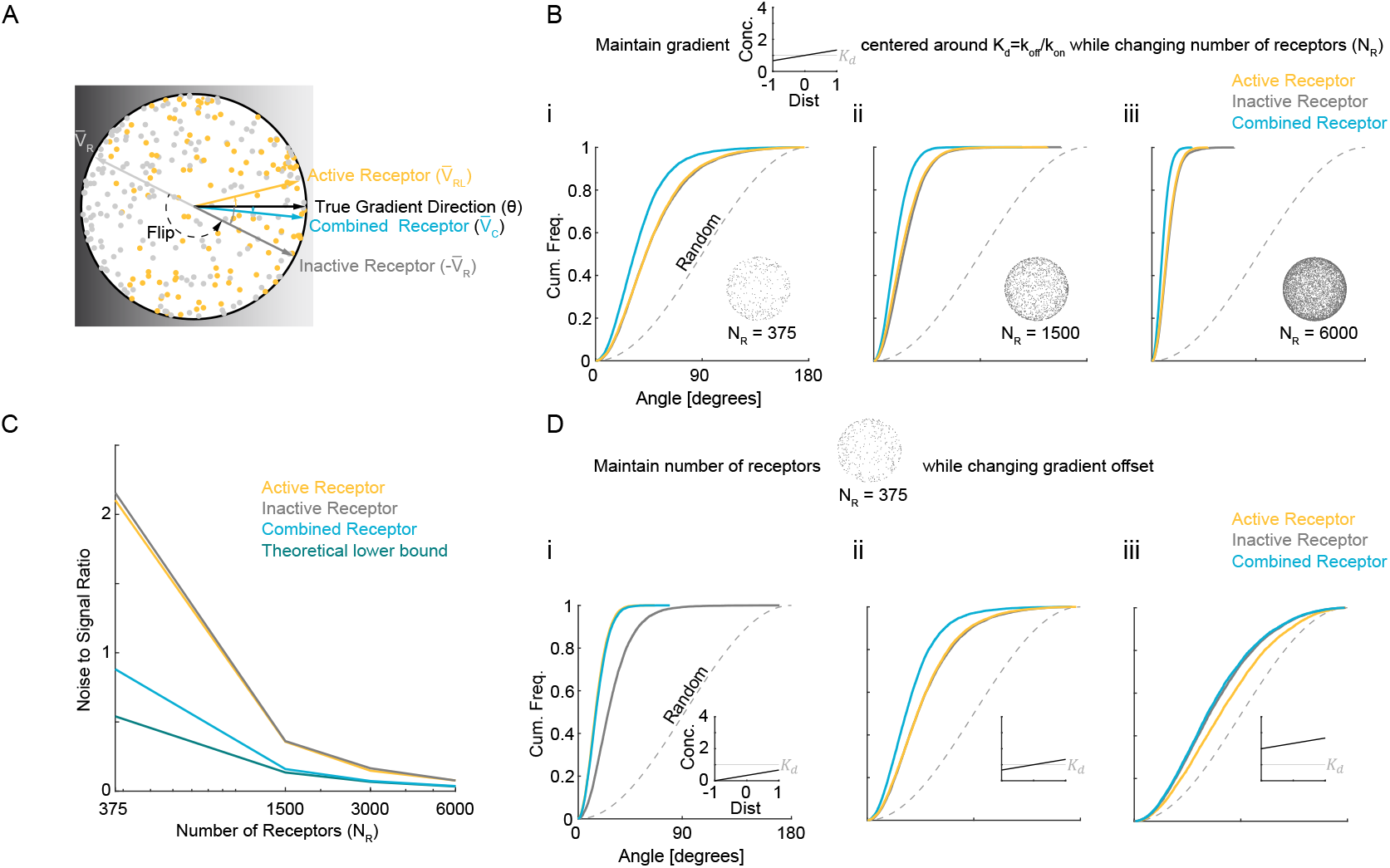
Estimating gradient direction using information from both active and inactive receptors. (**A**) Schematic depicting a spherical cell with receptors randomly distributed on its surface. Each receptor can be active (ligand-bound, yellow) or inactive (unbound, gray), and the gradient direction is inferred by summing receptor vectors and normalizing the sums to obtain unit vectors 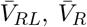and 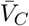. Note that 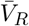 is flipped to obtain 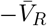 to estimate gradient direction. (**B**) Cumulative distribution functions (CDFs) of angular error for the estimates 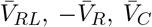, for three different receptor numbers (*N*_*R*_ = 375, 1500, 6000), where *K*_*d*_ = 6*nM* and the gradient is centered around *C*_0_ = *K*_*d*_ and ranges from 0.66 *K*_*d*_ to 1.33 *K*_*d*_ (i.e., *b/L* = 0.33). The dashed curve indicates the angular error for an estimate chosen uniformly at random over the cell. (**C**) Noise-to-signal ratio (NSR) comparing active-receptor sensing (yellow), inactive-receptor sensing (gray), and the combined-receptor estimate (blue) to the theoretical lower bound (dark green). Higher receptor counts reduce NSR, with the combined-receptor approach consistently outperforming either active- or inactive-only sensing. (**D**) Cumulative distribution functions (CDFs) of the angular error for the estimates 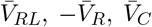 at fixed *N*_*R*_ = 375 but varying the gradient’s offset relative to *K*_*d*_: *C*_0_*/K*_*d*_ ∈ {1*/*3, 1, 3}. As the mean concentration shifts, active and inactive receptors each capture different portions of the ligand profile, and combining their information again improves direction inference.

The resulting active receptor vector, *V*_*RL*_, provides an estimate of the gradient direction pointing towards the highest ligand concentration. Comparing *V*_*RL*_ with the actual gradient direction provides an angular measure of how closely the cell’s guess matches reality. In principle, one could also compute the gradient direction from inactive receptors alone to produce a “best guess” vector *V*_*R*_ pointing towards the lowest ligand concentration. To combine information from both active and inactive receptors, we can calculate *V*_*C*_ = *V*_*RL*_ −*V*_*R*_ (Fig. 1A).

In what follows, we simulate a spherical cell in three dimensions, with *N*_*R*_ receptors randomly distributed over its surface. As receptor diffusion on the membrane is slow, especially in budding yeast (~ 0.0005*µm*^2^*/s*) [33, 36], we neglect it. Receptors can switch between active and inactive states at rates that depend on local ligand concentration *C*(*x*); in statistical equilibrium, the probability of a receptor being active at a given time is

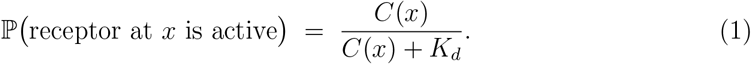

We expose the spherical cell to linear ligand gradients of the form

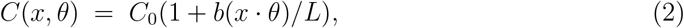

where *θ* is the direction vector, *C*_0_ is the mid-point (average) ligand concentration, and *x* is a point on the sphere of radius *L* centered at the origin. We first assume the gradient is centered around *K*_*d*_ (i.e., *C*_0_ = *K*_*d*_). Molecular noise from ligand binding/unbinding causes estimates of the gradient direction *θ* to fluctuate over time. We quantify directional accuracy by examining the empirical cumulative distribution functions for the angular deviation of 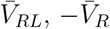, and 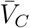 (the normalized summed vectors) relative to *θ* across many independent realizations (Fig. 1 B i-iii).

As 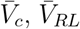, and 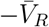 are each statistical estimators of gradient direction, we can compare them to the Cramér-Rao bound [37], which provides a rigorous lower bound on the variance of 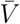 from the true direction *θ*, relative to the signal strength 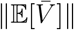. Accuracy of the direction estimate is quantified by a “noise-to-signal ratio” (NSR), and the statistical bound takes the form

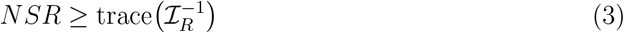

where ℐ_*R*_ is a Fisher information matrix that can be computed numerically. In the weak-signal regime (small slope, *b*) this bound is approximated by

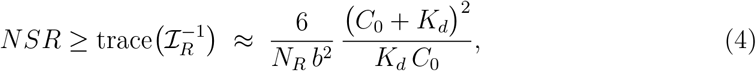

which shows that the minimal achievable variance in the estimated direction decreases proportionally to 1*/N*_*R*_ and depends inversely on *b*^2^, thereby highlighting the advantage of stronger gradients and higher receptor number (details of this bound and a derivation of (4) appear in the Supplementary Information). A related bound is derived in [32] for a different model.

A comparison with this theoretical lower bound (Equation 3) shows that the estimates 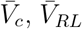, and 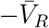 are indeed noisy, leaving ample room for improvement, especially at low *N*_*R*_ (Fig. 1 C). At high *N*_*R*_, the variance of 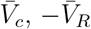, and 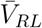 decrease along with the Cramér-Rao bound, which scales as 1*/N*_*R*_ (Fig. 1C). For *C*_0_ = *K*_*d*_, the variance of the combined estimate 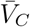 is roughly half that of either 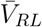 or 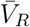, as expected from the 1*/N*_*R*_ scaling of the optimal variance and the fact that roughly half of the receptors are active in this regime. Effectively, 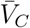 draws on twice the data that either 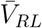 or 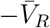 alone can provide, demonstrating that incorporating both ligand-bound and unbound receptors significantly enhances directional sensing.

To determine whether these conclusions extend to other gradients, we simulated a range of linear gradients covering different mean ligand concentrations (Fig. 1D). As the ligand concentration rises, saturation becomes more pronounced, so a larger fraction of receptors are active and the signal is diminished. Consistent with that expectation, *V*_*RL*_ performs increasingly poorly at higher concentrations (Figure 1D i-iii). Though −*V*_*R*_ also degrades at high concentrations, it does so differently from *V*_*RL*_. When the gradient is centered below *K*_*d*_, *V*_*RL*_ outperforms −*V*_*R*_ (Fig. 1D i); at *K*_*d*_, *V*_*RL*_ and −*V*_*R*_ perform similarly (Fig. 1D iii); above *K*_*d*_, −*V*_*R*_ becomes superior (Fig. 1D iii). This pattern suggests that at low ligand concentrations, the scarce active receptors provide more information than the many inactive ones, whereas at high ligand concentrations, the fewer inactive receptors carry more information. The combined vector *V*_*C*_ performs as well as or better than either *V*_*RL*_ or −*V*_*R*_ across the same tested range (Fig. 1 D i-iii). Hence, pooling signals from both active and inactive receptors broadens the range of gradients that can be decoded accurately.

### B. Ratiometric signaling biochemistry greatly enhances gradient estimation

Until now, we have imagined that a cell could apply some procedure to calculate *V*_*C*_ (and extract the direction 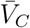), but we neglected the actual biochemical processes by which such calculations necessarily arise. Indeed, any downstream signaling step cannot preserve 100% of the information available at an earlier step, as noise in the interpretation mechanism will inevitably degrade the signal. For the ubiquitous and large family of G protein coupled receptors (GPCRs) that signal to downstream heterotrimeric G proteins, the biochemical mechanisms have been elucidated, and we consider here a simplified scheme in which a cell contains *N*_*G*_ G proteins that diffuse on the 2D surface of the spherical cell and can become activated or inactivated. Activation occurs with some probability upon collision between a diffusing inactive G protein and a ligand-bound receptor.

Following [33], we explore two different models of G protein inactivation (Fig. 2). In the *classical* model, inactivation occurs at a spatially uniform rate, 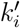, that is independent of receptor occupancy (Fig. 2A). In contrast, in the *ratiometric* model, inactivation occurs with some probability upon collision between a diffusing active G protein and an unbound receptor (Fig. 2B). We adjust the uniform inactivation rate of the classical model so that matched classical and ratiometric simulations have the same steady-state abundance of active G proteins, thereby permitting fair comparisons [33]. Note that unlike the ratiometric model, the classical model does not incorporate information about the spatial distribution of inactive receptors.

**FIG. 2.**
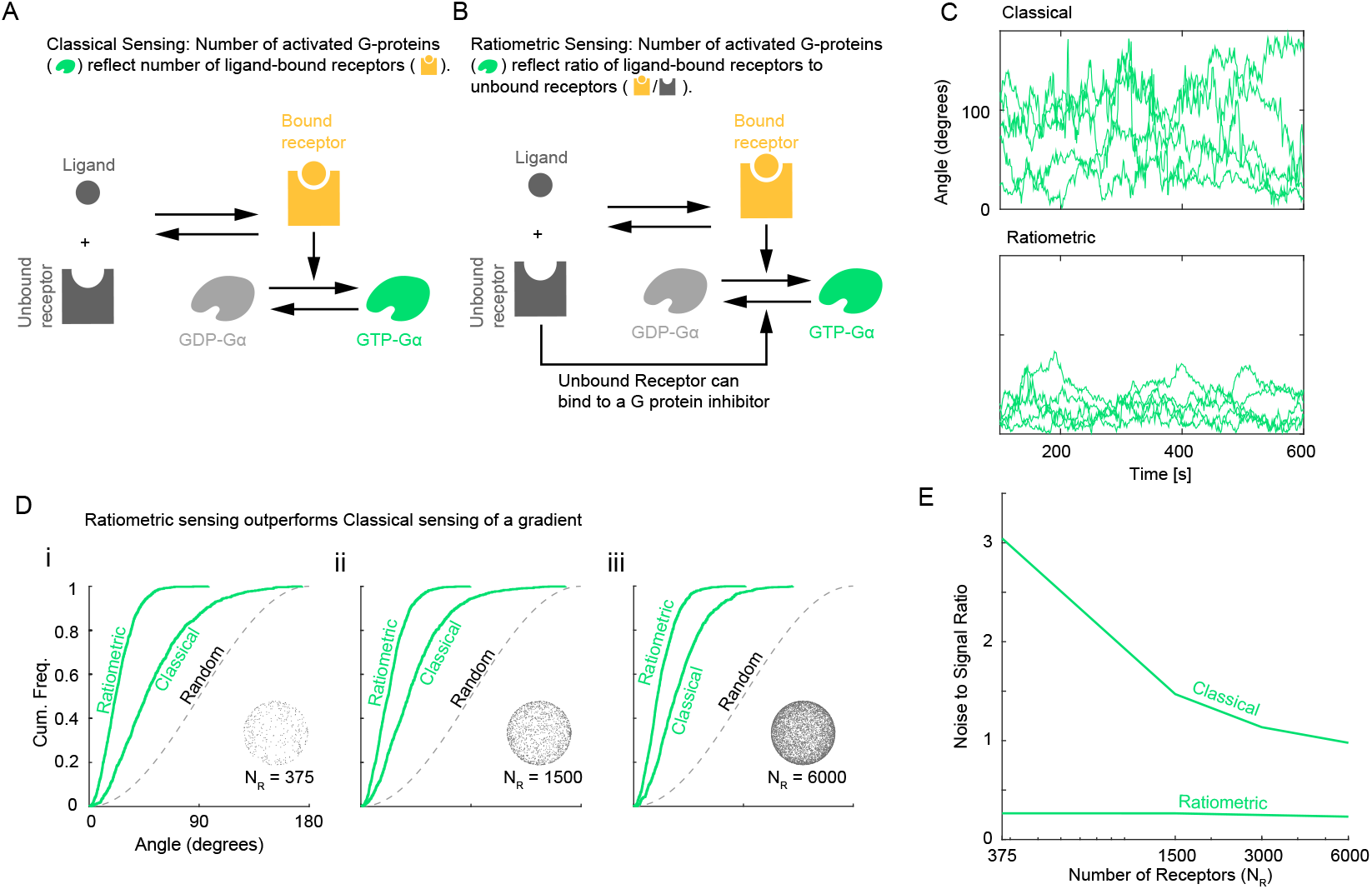
Ratiometric gradient sensing outperforms classical gradient sensing across a range of receptor numbers *N*_*R*_. (**A**) Schematic of reactions that constitute classical sensing. (**B**) Schematic of reactions that constitute ratiometric sensing. (**C**) 5 sample traces each of direction derived from G protein distribution over time for ratiometric and classical sensing strategies. *N*_*R*_ = 375, *N*_*G*_ = 2500, *D*_*G*_ = 0.002*µm*^2^*s*^−1^, *k*_*on*_ = 0.0167*s*^−1^*nM* ^−1^, *k*_*off*_ = 0.1*s*^−1^, *K*_*d*_ = 6*nM*. (**D**) Cumulative distribution functions of angular error comparing gradient sensing between ratiometric 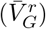 and classical 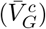 scenarios. *N*_*R*_ = [375, 3000, 6000] using the parameters in (C). (**E**) Noise to signal ratio of direction derived from G protein distribution compared between classical and ratiometric sensing strategies. *N*_*R*_ = [375, 1500, 3000, 6000] using the parameters in (C). For classical sensing, the G protein off-rate was assigned according to *N*_*R*_ ↦ *k*_*go*_ as [375, 1500, 3000, 6000] ↦ [0.0083, 0.0357, 0.0769, 0.1687]s^−1^.

In a ratiometric system, because ligand binding simultaneously increases the number of active receptors and decreases the number of inactive receptors, these opposing effects can amplify fluctuations due to molecular noise [38]. Indeed, [39] suggests that for a fixed ligand concentration, a cell’s measurement of ligand levels is noisier under a ratiometric (or “concerted”) mechanism than under the classical mechanism. Nevertheless, this need not imply that direction sensing becomes less reliable. For example, suppose (*F*_*r*_, *B*_*r*_) and (*F*_*c*_, *B*_*c*_) are pairs of random variables representing the active G protein concentration near the cell’s front (*F*) and back (*B*), under the ratiometric (*r*) or classical (*c*) mechanisms. It may be that *CV* (*F*_*c*_) < *CV* (*F*_*r*_) and *CV* (*B*_*c*_) < *CV* (*B*_*r*_), yet *F*_*r*_ − *B*_*r*_ > *F*_*c*_ − *B*_*c*_ (or *F*_*r*_*/B*_*r*_ > *F*_*c*_*/B*_*c*_) still holds with high probability. (*CV* denotes coefficient of variation.) In other words, although the “signal” at any given location may be noisier in the ratiometric mechanism, the overall direction information (e.g., the difference *F* − *B* or ratio *F/B*) can remain more robust compared to the classical system (see Section 5 and Fig. S6–S8 in the Supplement).

To compare direction sensing between classical and ratiometric systems, we calculated resultant vectors for active G proteins in a manner analogous to the previously discussed active receptor vector *V*_*RL*_ (Fig. 1A). We denote the resultant vectors of active G proteins under the ratiometric and classical models by 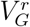 and 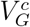, respectively, and their normalized directions by 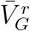 and 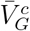. For ligand gradients near the receptor *K*_*d*_, the ratiometric model 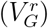 yields a significantly more accurate estimate of the gradient direction than the classical model 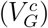 as demonstrated by temporal traces (Fig. 2C) (Movies S1-S6) and cumulative distribution functions (Fig. 2D) of 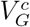 direction relative to gradient direction. Consequently, ratiometric sensing markedly enhances the accuracy of spatial gradient detection, even though it may sacrifice some accuracy in estimating the overall ligand concentration.

We next compared the performance of 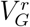 to that of the combined receptor vector *V*_*C*_. In simulations with many receptors (large *N*_*R*_), the receptor-based vector *V*_*C*_ performed better than 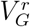 (Fig.3A). This was expected, as the smaller number of G proteins (*N*_*G*_ = 2500) cannot retain all directional information from the larger number of receptors (*N*_*R*_ = 6000). However, as *N*_*R*_ decreased, *V*_*C*_ became increasingly susceptible to stochastic noise. Remarkably, 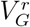 retained its performance at lower *N*_*R*_ values (Fig.3A, B), so that below some threshold in receptor count (approx. *N*_*R*_ < 1280), 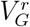 actually outperformed *V*_*C*_ (Fig. 3C). Even more unexpectedly, for sufficiently low *N*_*R*_ (approx. *N*_*R*_ < 800), 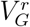 outperformed the accuracy limit given by the theoretical Cramér–Rao bound for *V*_*C*_ with the same number of receptors (Fig.3C). The Cramér-Rao bound (3) applies only to direction estimates based on instantaneous receptor data (i.e., *V*_*C*_, *V*_*R*_, *V*_*RL*_) and need not apply to 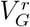 or 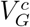; our finding shows that the accuracy of 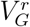 in this regime cannot be matched by any unbiased estimator based only on instantaneous receptors states at a single fixed time. While these simulations were carried out using custom code written in MATLAB 2024a, the surprising results were verified independently using the particle-based simulator SMOLDYN (Figures S1 and S2). Thus, ratiometric G protein signaling can continue to provide accurate directional information despite receptor scarcity.

**FIG. 3.**
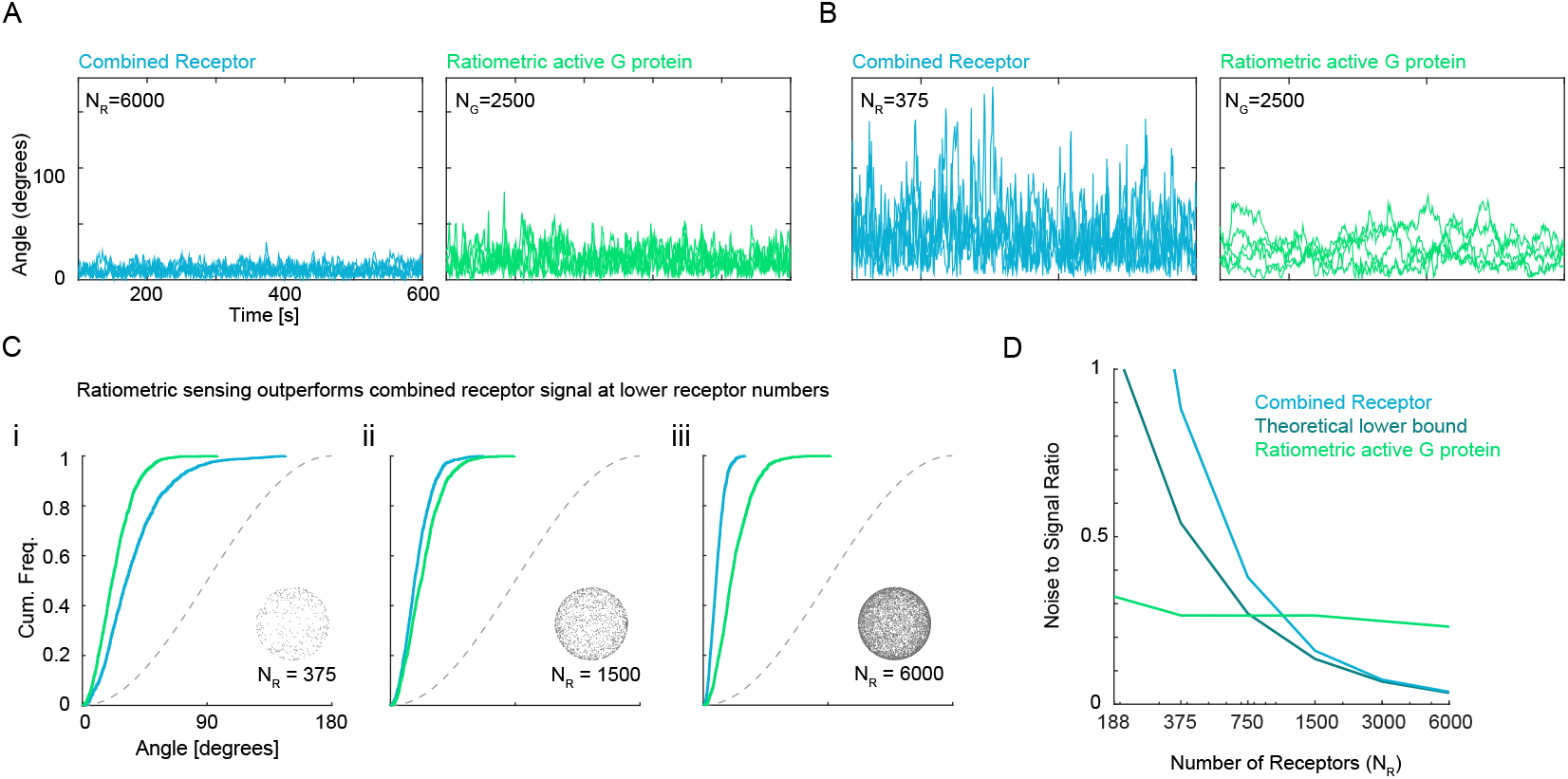
Ratiometric sensing allows G proteins to outperform instantaneous theoretical limit. (**A)** Temporal tracks of *V*_*C*_ and 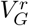 for *N*_*R*_ = 6000, *N*_*G*_ = 2500, *D*_*G*_ = 0.002*µm*^2^*s*^−1^,*k*_*on*_ = 0.0167*s*^−1^*nM* ^−1^,*k*_*off*_ = 0.1*s*^−1^, *K*_*d*_ = 6*nM*. (**B**) Temporal tracks of *V*_*C*_ and 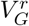 for *N*_*R*_ = 375 using parameters in (A). **(C)** Cumulative distribution functions of angular error comparing gradient sensing efficiency of active G protein in the ratiometric scenario to combined receptor. *N*_*R*_ = [375, 3000, 6000] using parameters in (A). (**D**) Comparison of noise-to-signal ratio with theoretical Cramér-Rao bound which applies to the estimate *V*_*C*_. For low values of *N*_*R*_, the noise-to-signal ratio for 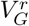 is below this theoretical bound, suggesting that 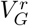 utilizes more information than is present in receptor states at a fixed time. *N*_*R*_ = [188, 375, 750, 1500, 3000, 6000] using parameters in (A)

### C. Ratiometric sensing’s enhanced gradient estimation arises from robust collective G protein memory

In all parameter regimes that we explored with our simulations, the ratiometric model 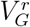 is more accurate in direction estimation than the classical model 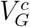. In some parameter regimes explored in our simulations, the ratiometric model 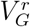 also outperforms both *V*_*RL*_ and *V*_*C*_, and it even performs better than the theoretical bound for estimation based on instantaneous receptor states (Fig. 3C). Here we explore the reasons for this superior performance of the ratiometric model.

The observation that 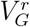 may perform better than both *V*_*RL*_ and *V*_*C*_, even better than the theoretical bound for estimation based on instantaneous receptor states, suggests that the distribution of active G proteins must carry more information than the instantaneous distribution of receptors alone. Because receptors and G proteins occupy binary states, the G protein population can only carry additional information if *N*_*G*_ > *N*_*R*_. Indeed, reducing the number of G proteins lowers the accuracy of 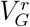 (Fig. 4A).

**FIG. 4.**
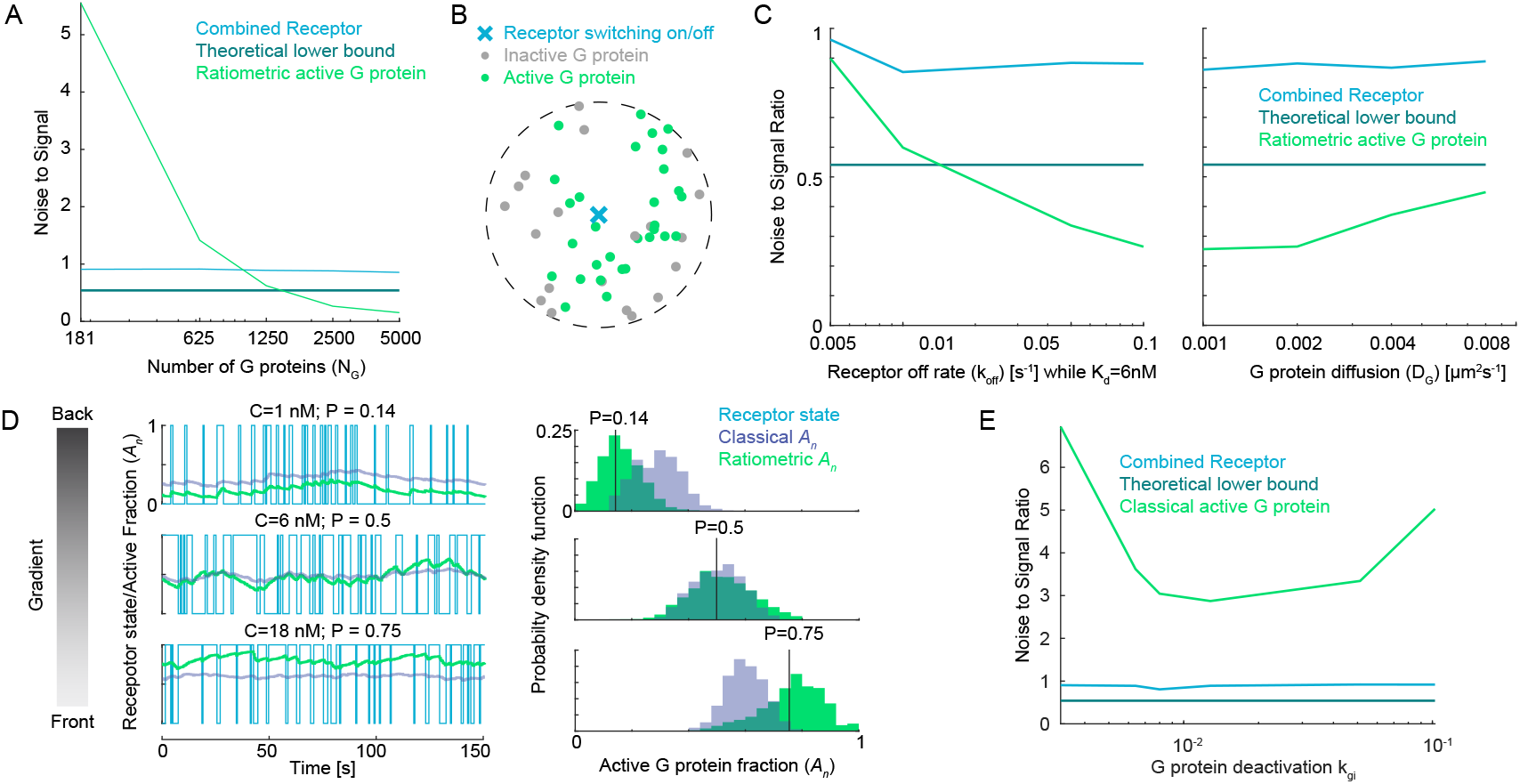
Ratiometric sensing enables G proteins to robustly generate a local average of past receptor states. (**A**) Noise-to-signal ratio for 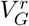, *V*_*C*_, and the theoretical lower bound as a function of the number of G proteins. *N*_*G*_ = [181, 625, 1250, 2500, 5000], *N*_*R*_ = 375, *D*_*G*_ = 0.002*µm*^2^*s*^−1^, *k*_*on*_ = 0.0167*s*^−1^*nM* ^−1^, *k*_*off*_ = 0.1*s*^−1^, *K*_*d*_ = 6*nM*. (**B**) Cartoon of a single receptor surrounded by G proteins that have recently inherited the receptor’s state. For ratio-metric signaling, G proteins can inherit state from ON or OFF receptors. For classical signaling, G proteins inherit state from ON receptors, but switch off at basal rate *k*_*gi*_.(**C**) Gradient sensing accuracy (NSR) as a function of receptor switching rate (left: *k*_*off*_ = [0.005, 0.01, 0.05, 0.1] *s*^−1^) and G protein diffusion constant (right: *D*_*G*_ = [0.001, 0.002, 0.004, 0.008] *µm*^2^*s*^−1^). *N*_*G*_ = 2500, *N*_*R*_ = 375. (**D**) Left: Temporal traces of the active G protein fraction for ratiometric and classical signaling at three positions along a gradient corresponding to different mean ligand concentrations: back (*C* = 1 *nM, p*_*R*_ = 0.14), middle (*C* = 6 *nM, p*_*R*_ = 0.5), and front (*C* = 18 *nM, p*_*R*_ = 0.75). The classical model is tuned to match ratiometric at *p*_*R*_ = 0.5. Right: Histograms of the active G protein fraction showing that the classical model deviates from *p*_*R*_ at the front and back, while the ratiometric model remains unbiased across all three regions. (**E**) Sweeping through different values of *k*_*gi*_ in the classical model shows optimal performance that does not cross instantaneous theoretical limits. *N*_*G*_ = 2500, *N*_*R*_ = 375. *k*_*gi*_ = [0.0032, 0.0064, 0.0080, 0.0128, 0.0512, 0.1024]

We propose that when *N*_*G*_ > *N*_*R*_, the collection of G proteins captures additional information by repeatedly sampling receptor states over time, thus storing past receptor states. The accuracy of the direction estimate 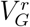 depends on the level of correlation in G protein states, which is governed by two key time scales. The first is the characteristic timescale of receptor state switching, λ^−1^, where

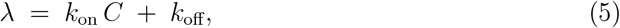

*k*_on_ and *k*_off_ are on/off rates of ligand binding, and *C* is the local ligand concentration. The second is the timescale of diffusive encounter, *τ*_dif_, which is the mean time between G protein-receptor collisions. This depends on the G protein diffusion constant (*D*_*G*_) and receptor number (*N*_*R*_):

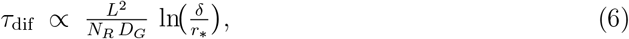

where *L* is the cell’s linear dimension (e.g., radius), *r*_∗_ is the receptor interaction radius, and *δ* is the typical receptor spacing. In the ratiometric model, it is the product *λτ*_dif_ that determines the degree to which the G protein states are correlated.

We consider a simplified model with *n* G proteins diffusing in the neighborhood of a single receptor that stochastically switches between active and inactive states, switching to active at rate *k*_on_*C* and to inactive at rate *k*_off_ (Fig.4B). As the receptor is switching states, the *n* G proteins encounter the receptor at random intervals with mean collision time *τ*_dif_. In the ratiometric version of this simplified model, G proteins activate upon encountering an active receptor but deactivate only upon collision with an inactive receptor. In this case, the mean fraction of active G proteins (*A*_*n*_) is the same as the fraction of time the receptor is on: 𝔼 [*A*_*n*_] = *p*_*R*_, but the variance of *A*_*n*_ changes with *τ*_dif_ and λ. A lower value of λ would lead to the receptor spending more time in a given state, causing states of G proteins to become more correlated. We derive an expression for variance of *A*_*n*_ (see Supplement):

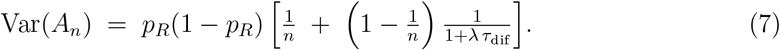

Thus, faster receptor switching (larger λ) or slower G protein diffusion (larger *τ*_dif_) raises *λτ*_dif_, which lowers Var(*A*_*n*_) and improves the accuracy of estimating *p*_*R*_. In Section 4 of the Supplement, we explain how this mechanism works when there are multiple receptors distributed across the cell surface, and we show how the timescales λ^−1^ and *τ*_dif_ affect the reliability of the direction estimate 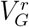. In our simulations, increasing λ (by increasing *k*_*off*_ while keeping *K*_*d*_ constant) leads to improved gradient sensing accuracy (lower NSR) (4C left), while increasing *D*_*G*_ leads to degraded accuracy (higher NSR) (4C right).

This analysis supports the striking observation that the noise-to-signal ratio of 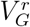 is relatively insensitive to *N*_*R*_ across a range of values of *N*_*R*_ that extends well below *N*_*G*_. Figure 2E, Figure 2C, and Figure 3D each demonstrate this insensitivity over a broad range. In Figure 3D, at low receptor number, 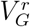 maintains a low noise-to-signal ratio—even below what is theoretically achievable for *V*_*C*_ at the same *N*_*R*_ (i.e., below the Cramér–Rao bound for *V*_*C*_). This occurs because historical sampling of multiple receptor states enables 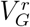 to encode more information than is available in instantaneous receptor states. In Figure 4A, as *N*_*G*_ increases (with *N*_*R*_ fixed), the estimate 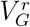 becomes more accurate than *V*_*C*_, the noise-to-signal ratio of 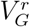 ultimately dropping below the Cramér–Rao bound for *V*_*C*_.

In principle, the classical model 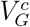 could also benefit from this time averaging, yet we observe that 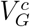 always performs worse than 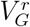. The reason for this difference in performance has to do with a dose-response bias that is inherent in the classical model. For concentration estimation (rather than gradient direction estimation), this bias was observed already in [35]. For the ratiometric model, each G protein’s state corresponds to the state of the receptor it most recently encountered. Consequently, the probability 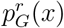 that a G protein at position *x* is active (in the ratiometric model) corresponds approximately to *p*_*R*_(*x*), the probability that a receptor at *x* is active. For the classical model, where a G protein’s state is reset to inactivate at rate *k*_*gi*_, the probability 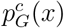 that a G protein at *x* is active depends on how many receptors it encountered since its last resetting event. This number of encounters depends on *k*_*gi*_ and on the diffusive time scale *τ*_dif_. In the supplement we derive the formula

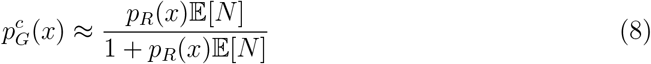

where 𝔼 [*N*] = *τ*_off_*/τ*_dif_ is the mean number of encounters, and 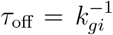. A consequence of this is that 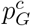 is not equal to *p*_*R*_ across the cell, and typically the gradient of 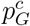 is significantly flatter than that of *p*_*R*_. Since the NSR for direction estimation is inversely proportional to the square of the ligand gradient (e.g., 1*/b*^2^ in (4)), this flattening of the gradient of 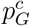relative to that of *p*_*R*_ implies a loss of accuracy in direction estimation for 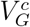.

To see how this bias effect may affect gradient sensing, we compared simulations of the ratiometric and classical versions of the simplified model described above. In the classical version, the *n* G proteins become activated upon collision with an active receptor and deactivate spontaneously at a baseline rate *k*_*gi*_. To compare these models, we examined the active G protein fraction (*A*_*n*_) at three positions across the cell corresponding to different ligand concentrations: back (low concentration, *p*_*R*_ = 0.14), middle (*p*_*R*_ = 0.5), and front (high concentration, *p*_*R*_ = 0.75) (Fig. 4D). At the middle position, where *p*_*R*_ = 0.5, we can tune *k*_*gi*_ so that the classical model matches the ratiometric model, with both accurately reflecting the receptor state. However, at the front and back, where receptors experience higher or lower ligand concentrations, the classical model with this fixed *k*_*gi*_ deviates from *p*_*R*_, while the ratiometric model maintains an unbiased estimate across all three regions without any parameter adjustment. Consequently, the ratiometric model produces a steeper gradient from front to back than the classical case.

This observation suggests the possibility of optimally tuning *k*_*gi*_ to improve gradient sensing in the classical model. We therefore swept through different values of *k*_*gi*_ in our full cellular gradient-sensing simulations (Fig. 4E). While we identified an optimal *k*_*gi*_ value that minimized the noise-to-signal ratio, this optimized classical model still performed worse than the combined receptor estimate *V*_*C*_ and failed to surpass the instantaneous theoretical bound, which is markedly inferior to the ratiometric model (Fig. 3D). Thus, ratiometric signaling enables robust collective memory through unbiased local averaging of receptor states, allowing cells to surpass the instantaneous theoretical limit for gradient detection.

### D. Responsiveness to changing gradients

In the preceding discussion, we assumed that ligand gradients were stable over time. However, this is not always the case: neutrophils chase motile bacteria, social amoebae collectively generate fruiting bodies by following dynamically shifting cAMP signals, and budding yeast track their mating partners as both cells grow [10, 40, 41]. While the robust collective memory encoded in G protein states by ratiometric signaling improves accuracy at steady state, it may slow responses to changing gradient directions.

To assess how different parameters affect a cell’s ability to adapt to changing gradients, we performed simulations in which we allowed 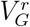 to reach steady state in a gradient and then instantaneously reversed the gradient direction by 180 degrees (Fig. 5A). We quantified responsiveness by measuring the time required for 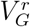 to cross the 90-degree threshold toward the new gradient direction (Fig. 5B).

**FIG. 5.**
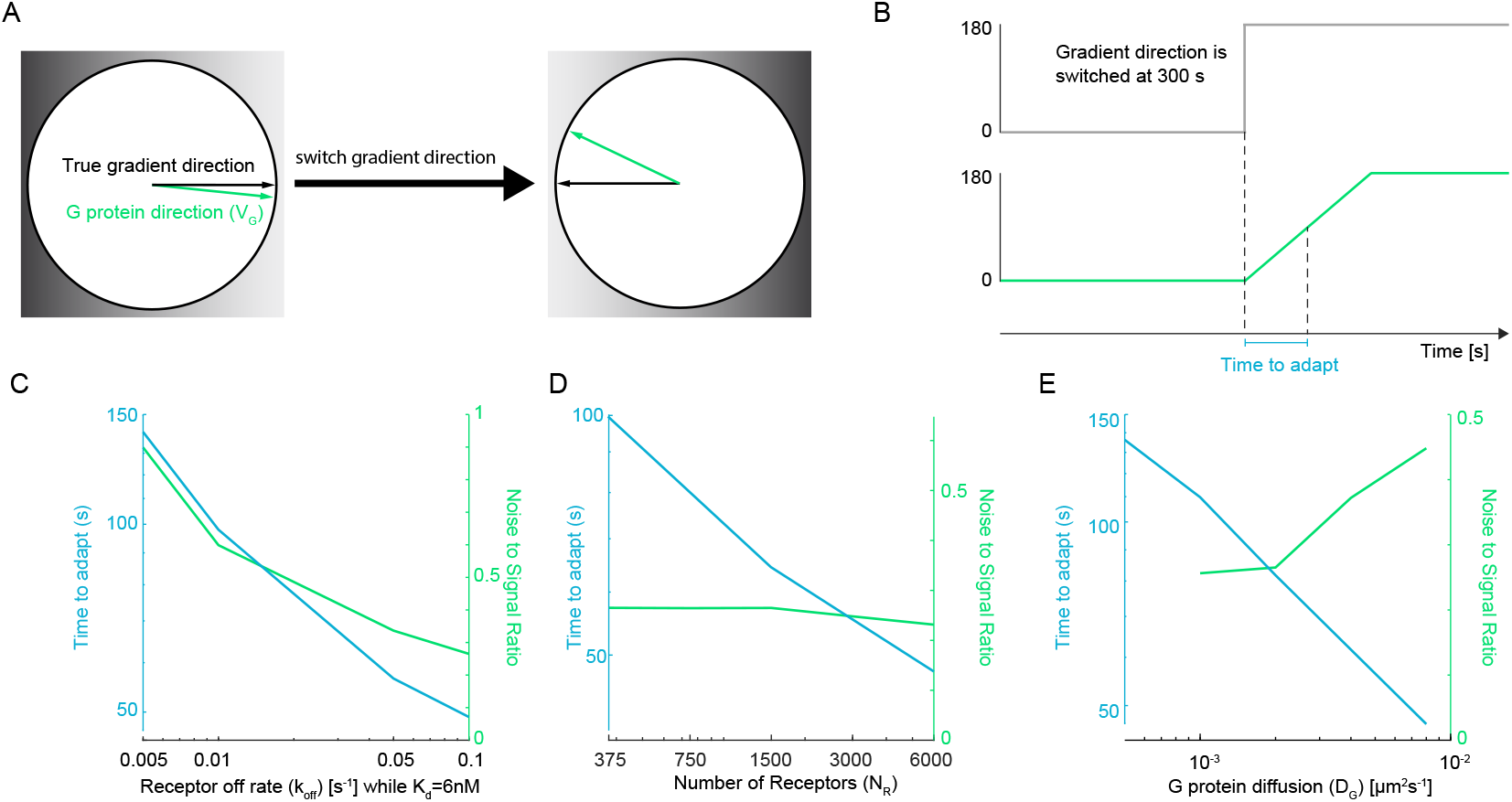
Adaptation to changing gradients. Default simulation parameters: *N*_*R*_ = 375, *N*_*G*_ = 2500, *k*_off_ = 0.1 s^−1^, *K*_*d*_ = 6 nM, and *D*_*G*_ = 0.002 *µ*m^2^s^−1^. (**A**) Cartoon demonstrating simulation where the true gradient direction (black arrow) is flipped by 180°. The G protein direction vector *V*_*G*_ (green arrow) eventually re-orients to match the new gradient. (**B**) Cartoon demonstrating how adaptation time of gradient estimate by G proteins is calculated. Top: The environmental gradient direction is switched at *t* = 300 s from 0° to 180°. Bottom: The direction of the averaged trace derived from 1000 G protein vectors 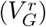 lags before reaching the new steady state. The time to adapt is defined as the interval required to cross 90° toward the new directional equilibrium. (**C–E**) Parameter sensitivity of adaptation speed and sensing accuracy. Blue curves (left y-axis) indicate time to adapt; green curves (right y-axis) indicate the steady-state noise-to-signal ratio (NSR). (**C**) Sweeping receptor off-rate (*k*_off_) from 0.005 to 0.1 s^−1^ shows that faster receptor kinetics improve both adaptation speed and directional accuracy. (**D**) Sweeping the number of receptors (*N*_*R*_) from 375 to 6000 shows that higher receptor density significantly reduces the time to adapt while maintaining a relatively robust NSR. (**E**) Sweeping G protein diffusion (*D*_*G*_) from 10^−3^ to 10^−2^ *µ*m^2^s^−1^ shows that faster diffusion accelerates adaptation but leads to a higher NSR, highlighting a fundamental trade-off between response speed and spatial precision.

We first examined the effect of receptor switching kinetics by varying both ligand off-rate *k*_off_ and on-rate *k*_*on*_ while maintaining a constant *K*_*d*_ (Fig. 5C). While *k*_on_ is physically capped by the rate of diffusional encounters, our parameters remain biologically plausible; at *K*_*d*_ = 6 nM, the maximum *k*_off_ = 0.1 s^−1^ corresponds to *k*_on_ ≈ 1.67×10^7^ M^−1^s^−1^, which is well within the Smoluchowski limit. Increasing *k*_off_ from 0.005 to 0.1 s^−1^ reduced both the time to adapt and the steady-state noise-to-signal ratio. These dual benefits can be understood through the receptor switching rate λ (Equation 5): increasing *k*_off_ increases λ, which directly accelerates receptor state transitions and reduces the time to achieve statistical equilibrium after the gradient direction is reversed. Simultaneously, higher λ increases the product *λτ*_dif_; as we have discussed, this reduces correlation in downstream G protein states, and thus improves steady-state directional accuracy.

Increasing receptor density also accelerated adaptation to new gradient directions (Fig. 5D). Higher *N*_*R*_ reduces the mean time between G protein–receptor encounters (*τ*_dif_), allowing G proteins to update their states more frequently and thereby adapt more quickly to the reversed gradient. As we have already noted, the noise-to-signal ratio of 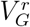 (at steady state) may be relatively insensitive to changes in *N*_*R*_ over a certain range, depending on *N*_*G*_ (e.g., as *N*_*R*_ increased from 375 to 6000 in Fig. 5D).

In contrast to receptor kinetics and density, increasing the G protein diffusion constant *D*_*G*_ presented a trade-off between adaptation speed and steady-state accuracy (Fig. 5E). Faster diffusion reduces *τ*_dif_, enabling more frequent G protein–receptor encounters and thereby accelerating adaptation to the new gradient direction. However, unlike increasing *N*_*R*_, faster diffusion of G protein increases correlation in G protein states and increases the noise-to-signal ratio of 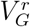 at steady state.

Together, these results reveal distinct strategies by which cells could tune their responsiveness to changing gradients. Faster receptor kinetics provide the most favorable outcome, simultaneously enhancing both steady-state accuracy and adaptation speed. Increased receptor density primarily accelerates adaptation with minimal impact on steady-state performance. Higher G protein diffusion offers faster adaptation but compromises spatial precision—a trade-off that cells may exploit depending on whether their environment demands accurate gradient tracking or rapid reorientation.

## III. DISCUSSION

### A. Ratiometric signaling in the budding yeast system

Gradient decoding is most challenging when cells are small, gradients are shallow, and chemical concentrations are low, but yeast cells find mates despite encountering all of these challenges. An effective strategy to extract signals from noisy measurements would be to integrate concentration measurements over time, allowing time averaging to reduce the noise (fluctuations) without affecting the signal (gradient). However, once a chemical has bound to a cell-surface receptor, diffusion of the receptor-ligand complex would degrade the information about the location where the binding took place. Such blurring can limit the interval over which time-averaging is effective for directional sensing[35]. This is particularly problematic for systems like yeast pheromone sensing, where ligand-receptor complexes can persist for several minutes. Interestingly, yeast cells severely limit receptor diffusion (*D* < 0.0005 *µm*^2^*/s*), perhaps mitigating this problem [33]. However, with very slow receptor diffusion, receptors often accumulate non-uniformly on one side of the cell [33, 42–44]. Non-uniform receptor distributions in turn can lead to spatial patterns of ligand-receptor complexes that do not accurately reflect the spatial distribution of ligands in the environment.

Yeast cells appear to use a ratiometric sensing strategy to compensate for non-uniform receptor distributions [33]. Ligand binding to receptors promotes activation of G proteins that are then turned off by RGS proteins (Sst2 in yeast). In classical signaling models, G protein inactivation occurs in a receptor-independent manner, but in yeast cells the RGS protein is bound to inactive receptors [42]. Thus, whereas active receptors promote signaling, inactive receptors inhibit signaling, and the net signal reflects the ratio of active to inactive receptors [34]. If receptors are concentrated in some region of the cell surface, both active and inactive receptors may be locally enriched, so ratiometric sensing can allow the cell to accurately infer the local ligand distribution by compensating for the local receptor abundance. Interestingly, we find that, in addition to extracting information from both active and inactive receptors, ratiometric sensing could also better enable a cell to interpret temporally averaged receptor states over time.

### B. Noise in concentration measurement from ratiometric signaling

In principle, ratiometric signaling could be a widely applicable strategy to improve the detection of spatial gradients, especially when gradient concentrations are near the ligand-receptor *K*_*d*_. It seems intuitively plausible that a cell capable of using information about the spatial distributions of inactive as well as active receptors would have a firmer basis to infer the spatial distribution of ligand than a cell whose only information comes from the active receptors. However, as discussed in the Supplement and shown in [38, 39], ratiometric signaling can be less accurate at sensing absolute concentration. Nevertheless, we find that despite being worse at sensing ligand concentration, the ratiometric strategy provides more accurate gradient sensing by enabling acquisition of information from both active and inactive receptors as well as locally averaging receptor signals over time.

We show that ratiometric signaling outperforms classical signaling, even at very low receptor numbers when less than 0.02% of the cell surface is covered in receptors. Notably, these conditions resemble scenarios where cells initially populate their membranes with specific receptors—such as in lymphocytes like B cells, which upregulate receptors upon transitioning between germinal center zones; and in various developmental contexts where cells switch states and begin expressing new receptors [45, 46]. In such scenarios, ratiometric signaling could enable reliable gradient detection with lower receptor density, all while resisting misdirection by local fluctuations in receptor localization or ligand availability.

### C. Mechanism underlying gradient detection accuracy by ratiometric signaling

One of our most intriguing observations was that ratiometric signaling could outperform the theoretical bound for gradient direction estimates that use only receptor snapshots at a single moment in time. This surprising advantage arises because downstream molecules—specifically, diffusing G proteins—act as a form of collective molecular memory. As each G protein updates its activity state only upon encountering a receptor, it encodes receptor information from an earlier time point. A population of G proteins that do this for both active and inactive receptors can smooth out short-lived fluctuations and enhance the fidelity of direction estimates. In effect, G proteins collectively time-average receptor states, allowing the cell to extract more robust spatial information than would be possible from instantaneous receptor states alone.

In ratiometric signaling, two parameters critically balance directional accuracy and adaptive speed: the switching rate (λ) of receptor-ligand binding and the diffusion coefficient (*D*_*G*_) of G proteins. Increasing λ enhances collective local averaging by G proteins (Fig. 4E (left) and 5C), improving steady-state gradient detection, and promotes faster adaptation to shifting gradients (Fig. 5C). In contrast, raising *D*_*G*_ weakens local averaging and diminishes steady state-gradient detection (Fig. 4E (right) and 5E). But higher *D*_*G*_ accelerates adaptation to changing signals, as more frequent receptor–G protein collisions swiftly update G protein states in response to new external conditions (Fig. 5C). Together, these trade-offs highlight how ratiometric signaling can be tuned to optimize either accuracy or speed, as required by the biological context.

### D. Broader relevance of ratiometric signaling in responding to directional cues

It is not known whether receptors other than GPCRs can engage ratiometric signaling pathways. However, the principle underlying the averaging of upstream signals by more abundant downstream components may extend beyond receptor signaling, particularly in systems where both active and inactive states of upstream proteins convey meaningful information. In MAPK cascades, downstream kinases often exceed the abundance of their upstream regulators, exemplified by Raf in mammalian ERK pathways [47], B-Raf in cancer signaling contexts [48], and Ste11 in yeast mating responses [49]. These systems possess features that suggest the potential for ratiometric measurement-driven averaging to enhance accuracy and robustness.

Overall, our work suggests that ratiometric signaling mechanisms provide a powerful framework for cells to interpret directional receptor information and mitigate noise. The ability to sample both active and inactive states, combined with downstream emergent molecular memory, enables more accurate direction finding than that achieved by classical signaling mechanisms. These results offer testable predictions for experimental systems and may help clarify why some cells have a higher abundance of downstream proteins in signaling cascades.

## IV. ACKNOWLEDGMENTS

This work was supported by NIH/NIGMS grant R35GM122488 to D.J.L. D.G. was supported by the Alfred P. Sloan Foundation’s Matter-to-Life program.

## V. DATA AVAILABILITY STATEMENT

The data and simulation code that support the findings of this study are openly available in the following GitHub repository: https://github.com/DebrajGhose/ratiometric-sensing-robust.

## Appendix A: Simulation Framework

The simulation models the interactions between receptors and G proteins on a cell surface, taken to be a sphere 𝒮 in three dimensions. The simulation focuses on their activation and inactivation dynamics under a pheromone gradient.

On the surface of the sphere, receptors are positioned randomly; their locations are fixed and do not change over time. Receptors also switch state randomly and independently of each other; they can be either ligand-bound (active) or unbound (inactive). The rate of activation depends on the local ligand concentration, which is taken to be time-independent.

The G proteins are also positioned randomly over the surface of the sphere, independently and uniformly distributed. Unlike the receptors, however, each G protein performs a diffusive random walk along the surface of 𝒮. G proteins in the simulation can exist in either an active or inactive state. The state of a G protein is influenced by its interaction with nearby receptors and its inherent deactivation rate. G proteins switch state according to either the classical model or the ratiometric model. In both models, a G protein becomes activated upon encountering an active receptor. The difference in the two models lies in the mechanism for deactivation. In the classical model, a G protein becomes deactivated randomly at the inherent deactivation rate. In the ratiometric model, a G protein becomes deactivated by encountering an inactive receptor.

We use the following nomenclature:

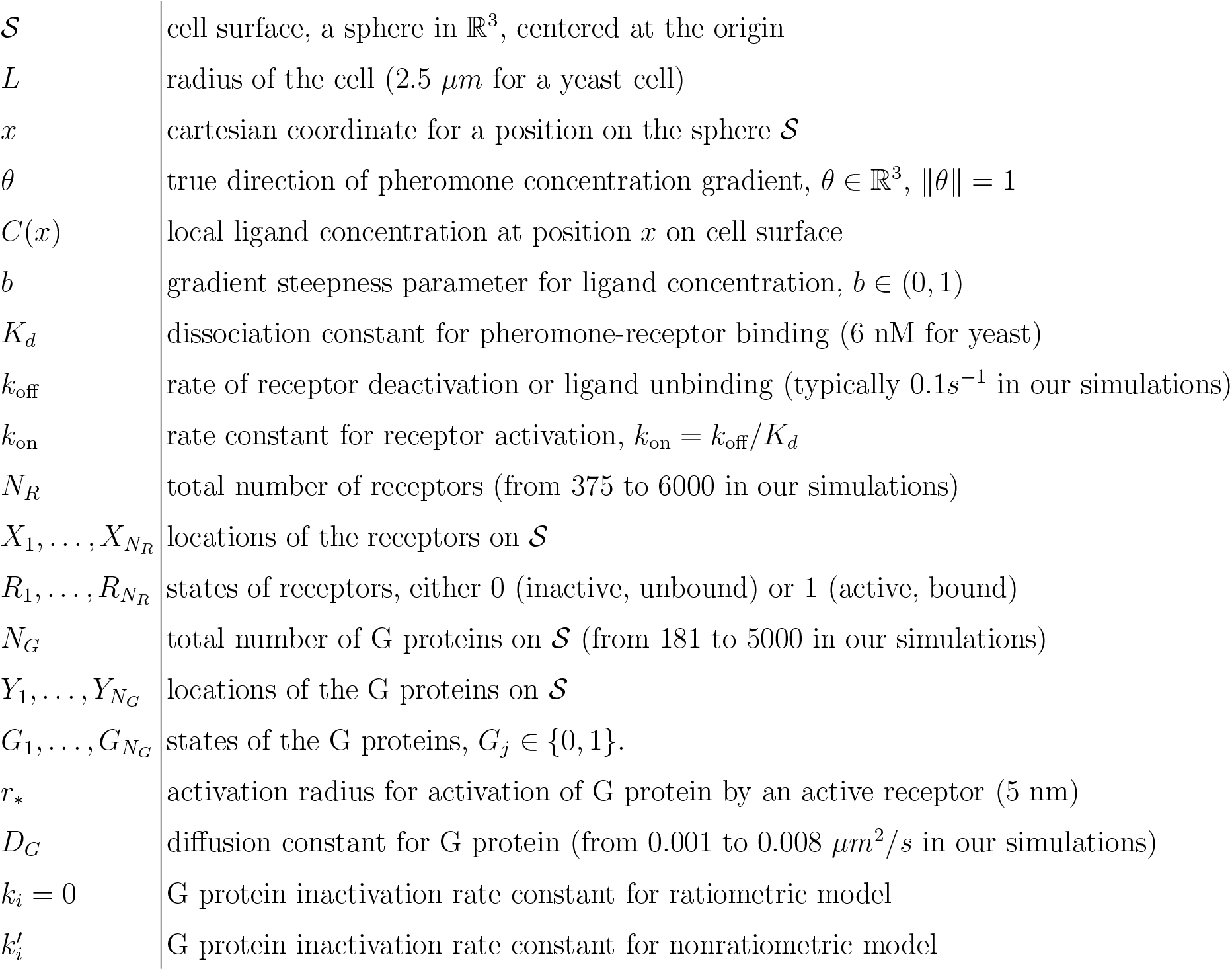

### 1. Receptor dynamics

The receptor configuration at time *t* is characterized by the pairs 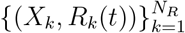. The positions 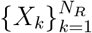 are sampled independently and uniformly on 𝒮. The receptor positions are assumed to not change with time (we neglect receptor diffusion). Given a receptor at position *X*_*k*_ = *x*, the receptor state *R*_*k*_(*t*) ∈ {0, 1} changes as a Markov process with rates:

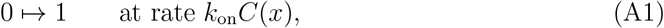

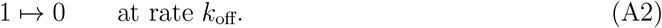

The receptor states are initialized according to the stationary distribution. Therefore, at any time, the probability that a receptor at location *X*_*k*_ = *x* is active is:

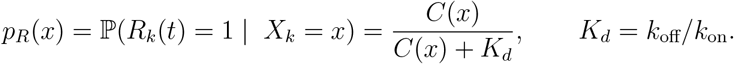

The ligand concentration depends on the direction vector *θ*, the true gradient direction. We will write *C* = *C*(*x, θ*) to emphasize this dependence on *θ*. We take *C* to have the linear form

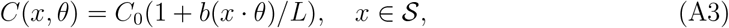

where *C*_0_ > 0 and *b* ∈ (0, 1), so that (1 − *b*)*C*_0_ ≤ *C*(*x, θ*) ≤ (1 + *b*)*C*_0_. Consequently, the probability *p*_*R*_(*x*) also depends on *θ*, and we emphasize this by writing *p*_*R*_(*x, θ*).

As we will explain later, the rate parameter

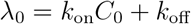

plays an important role in the model; 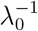 has the units of (*time*) and we refer to 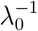 as the receptor turnover time.

We consider values of *N*_*R*_ in the range 375 to 6000 in our simulations. For a cell radius *L* = 2.5*µm* and receptor activation radius *r*_∗_ = 5*nm*, the fraction of area covered by receptor is in the range of 0.0375% to 0.6%.

#### 2. G protein dynamics

The G proteins are initialized independently, uniformly at random on the sphere, at positions 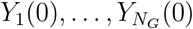. These positions diffuse over 𝒮 in time, independently and with diffusion constant *D*_*G*_; statistically, their joint spatial distribution is invariant in time. An inactive G protein (state *G*_*j*_(*t*) = 0) at location *Y*_*j*_(*t*) is activated upon interaction with an active receptor. Such activation occurs when *Y*_*j*_(*t*) diffuses within radius *r*_∗_ of an active receptor. The simulation uses a spatial partitioning algorithm (KD-tree) to identify, at each time step, G proteins within a certain reaction radius of any receptor.

Once active (*G*_*j*_ = 1), the mechanism for de-activation of G protein is either the ratio-metric model or the classical model. In the classical model, an active G protein deactivates at a fixed rate 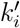, the inherent deactivation rate, but an active G protein is unaffected by encountering inactive receptors. In the ratiometric model, however, an active G protein deactivates upon diffusing within radius *r*_∗_ of an inactive receptor and there is no basal deactivation (*k*_*i*_ = 0 in the ratiometric model). To compare the ratiometric and classical models, we choose 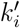 so that at steady state, the mean fraction of active G protein is the same as that for the ratiometric model. In particular, we adjust the parameter 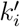 for each ligand concentration profile to maintain this comparison.

Depending on the random receptor configuration, it is possible for a G protein to be within the reaction radius of two or more receptors simultaneously. For the ratiometric model, when a G protein is near multiple receptors simultaneously the state (at each time step) of the G protein is matched that of a neighboring receptor chosen uniformly at random (from among those receptors within the reaction radius). Hence, if the majority of nearby receptors are active, the probability increases for the G protein to become active. Conversely, if most nearby receptors are inactive, the likelihood increases for the G protein to become inactive. This dynamic represents a ratiometric response of the G protein to the local receptor state.

#### 3. Simulation methodology and verification

Simulations were performed using custom solvers in MATLAB (v2021b to v2024a) on a Linux-based computing system (Longleaf cluster at UNC Chapel Hill). Simulations were initialized by randomly and uniformly distributing receptor and G protein populations on the surface of a spherical cell. Receptor positions were kept stationary while G proteins performed random walks (Brownian motion) across the cell’s surface with a fixed time step and diffusion constant. Receptor activation followed a first-order reaction, with an activation rate determined by the local pheromone concentration prescribed along the cell’s x-axis to establish a gradient (Equation (A2) and A 1). Activation and deactivation events were implemented via Monte Carlo sampling. In the classical receptor model, G proteins became active upon encountering active receptors within a specified reaction distance and deactivated at a constant rate. In the ratiometric model, G proteins were activated upon encountering active receptors within the same reaction distance and deactivated upon encountering inactive receptors. In the presence of multiple receptors, the ratio of active to inactive receptors governed the probability of G-protein activation or deactivation. For each set of parameters, 1000 simulations were run, tracking molecular states, positions, and time-resolved changes in receptor and G-protein activation. All MATLAB code for these simulations is available at: https://github.com/DebrajGhose/ratiometric-sensing-robust

To validate our solver, MATLAB simulations from Figures 1, 2D and 3C were replicated in Smoldyn (v2.71) with continuous space and discretized time intervals on a Linux-based computing system (Longleaf cluster at UNC Chapel Hill). Periodic boundary conditions were applied in both spatial directions. Molecules were modeled as point particles without volumes. Brownian motion of the molecules was simulated using the Euler-Maruyama method. We used a time step of 0.005 secs and recorded the coordinates of receptors and G proteins every 10 secs over 20000 secs. The simulation methods used were consistent with the customized code, except for the setup of the pheromone gradient. As Smoldyn does not support continuously spatially dependent concentration, the cell sphere was divided into 11 vertical compartments along the gradient direction, with each compartment assigned a local pheromone concentration defined as:

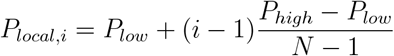

where *P*_*low*_ and *P*_*high*_ represent the lowest and highest pheromone concentrations, respectively. *i* is the compartment index. *N* = 11 is the total number of compartments. Accordingly, we modeled the receptor activation as a first-order reaction with a local activation rate in the *i*th compartment defined as:

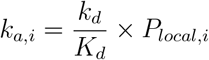

where *k*_*d*_ = 0.1*s*^−1^ is the dissociation rate between the receptor and the ligand.

All code for Smoldyn simulations are available at: https://github.com/DebrajGhose/ratiometricsensing-robust Figure 6 and 7 show that Smoldyn simulations produce similar results to those of our custom code.

**FIG. 6.**
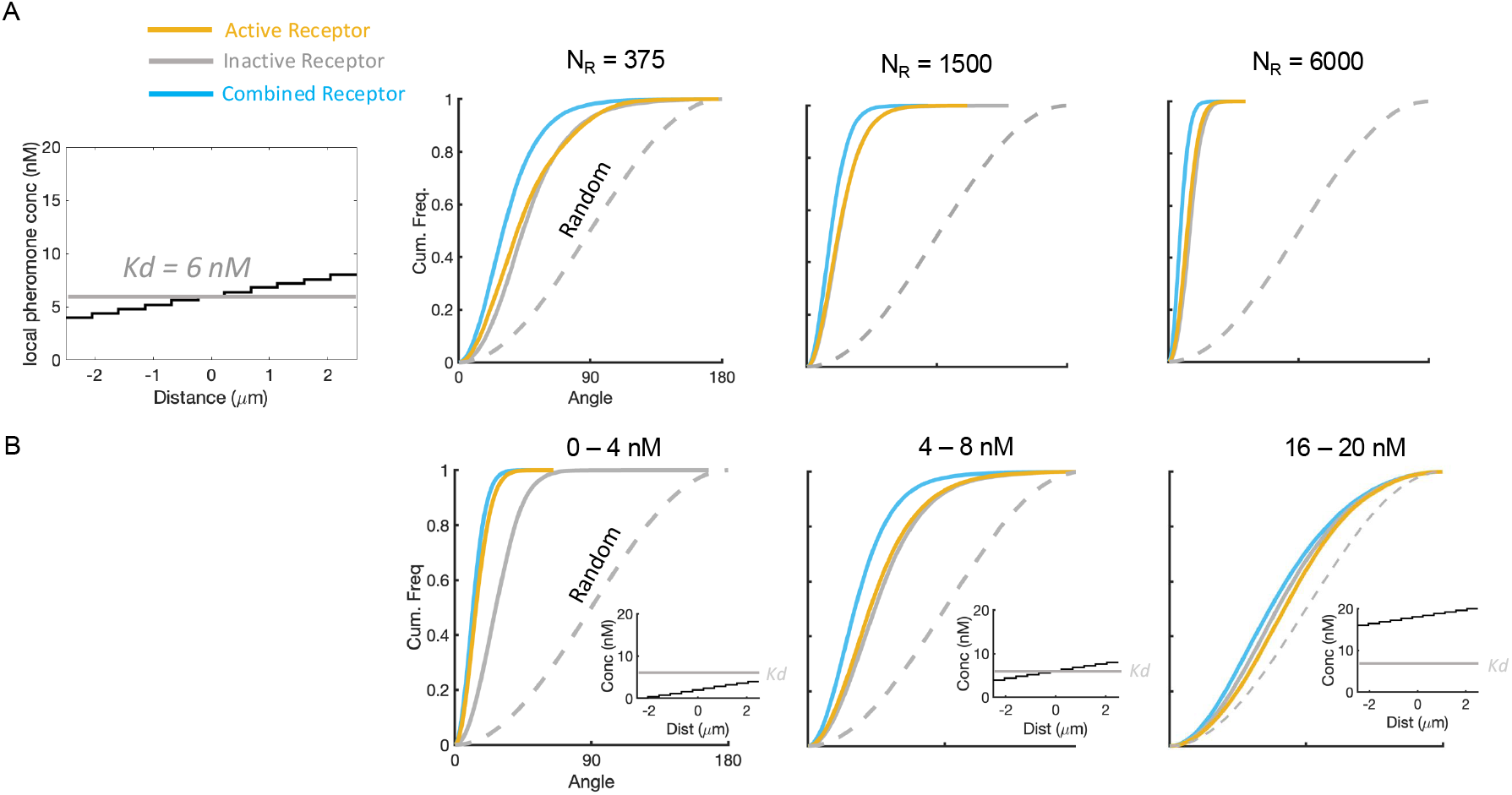
Replicating results in comparison of receptor signals using Smoldyn. **A)** Angular dis-tribution of receptors with the same pheromone gradient centered around *K*_*d*_ but under different receptor number. Corresponds to Fig. 1B. *n* = 10 cells. **B)** Angular distribution of receptors with the same receptor number but under different heights of pheromone gradients. Corresponds to Fig. 1D. *n* = 50 cells.

**FIG. 7.**
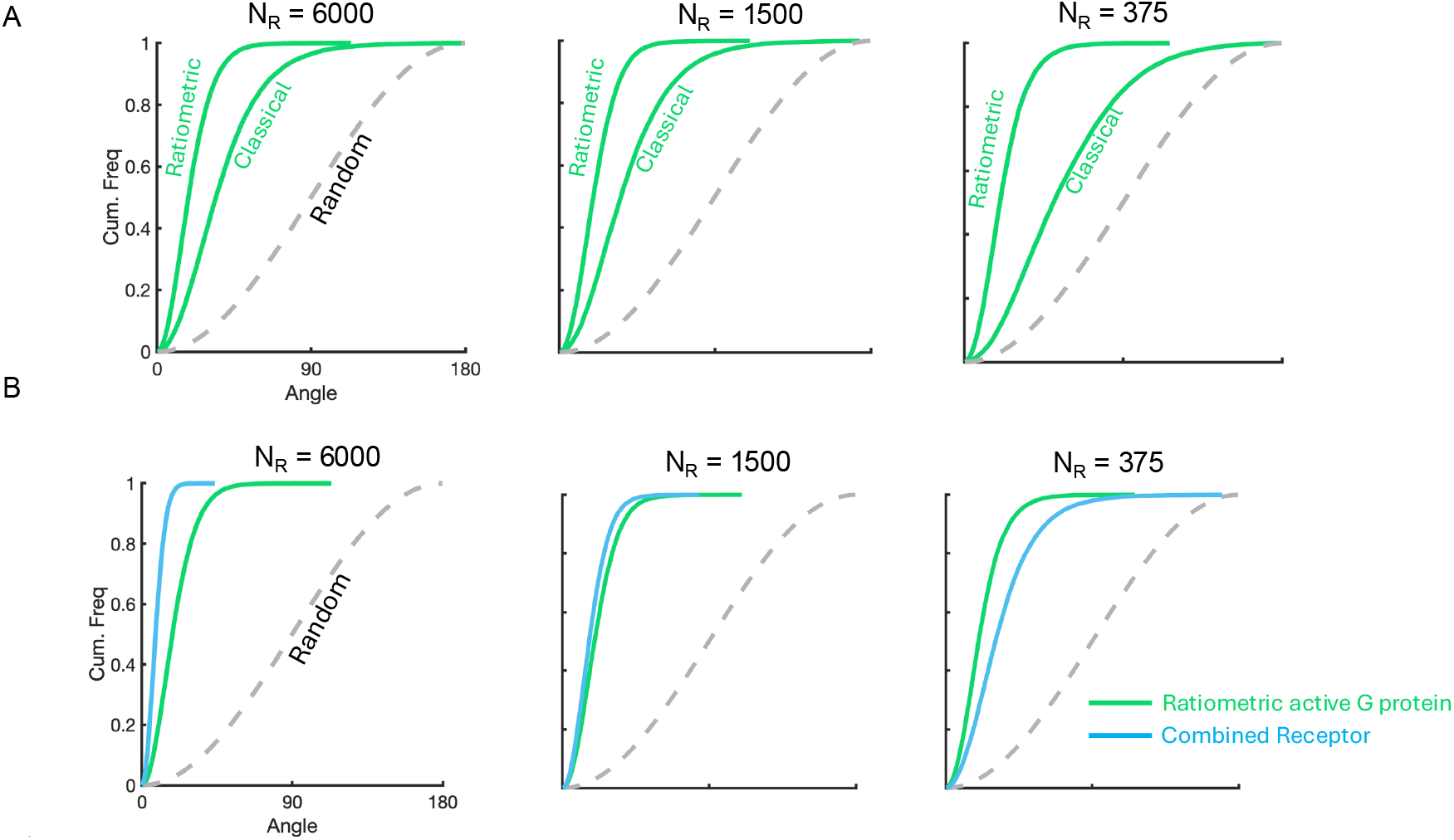
Replicating results in comparison between ratiometric and classical active G protein signal and combined receptor signal using Smoldyn. **A)** Angular distributions comparing gradient sensing between ratiometric and classical active G proteins. Corresponds to Fig. 2D. *n* = 10 cells. **B)** Angular distributions comparing gradient sensing between ratiometric active G proteins and combined receptors. Corresponds to Fig. 3C. *n* = 10 cells.

## Appendix B: Estimation model

At a given time *t*, we use the receptor data and the G protein data to compute various estimates of the direction parameter *θ*, treated as an unknown quantity. The receptor data at a fixed time are random variables 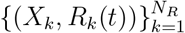. These variables are pairwise independent, and their distribution depends on the parameter *θ*. Thus, the receptor data may be regarded as *N*_*R*_ independent points in 𝒮 × {0, 1}, and their joint density on 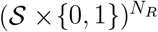 is

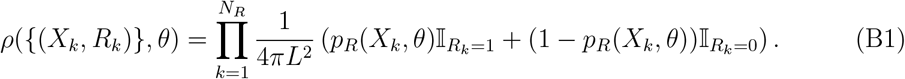

The factor 4*πL*^2^ is the surface area of 𝒮. The terms 𝕀_*R*=1_ and 𝕀_*R*=0_ are indicators of the events *R* = 1 and *R* = 0, respectively.

Using active receptor data, we define

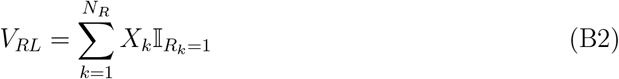

where 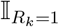 is the indicator of the event *R*_*k*_ = 1. Thus, *V*_*RL*_ is the sum of the positions of active receptors only. Similarly, using inactive receptors, we define

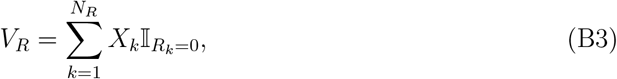

and we regard the reflected vector (−*V*_*R*_) as an estimate of *θ*, since the gradient of inactive receptor will typically be in a direction anti-aligned with *θ*. The combined receptor estimate is

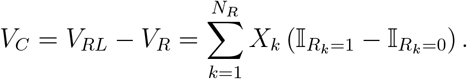

These *V*_*RL*_, −*V*_*R*_, and *V*_*C*_ are vectors in ℝ^3^. From these vectors, direction information is computed by normalizing

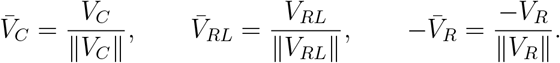

Each of these direction estimates is a random unit vector, depending on time (because the receptor states change in time). To compute the distribution of these estimates, we obtain independent samples by re-initializing independently all of the receptor positions and states.

Using the G protein data, we compute the vector

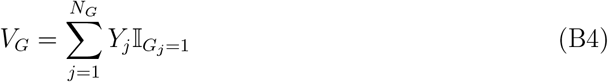

which is the sum of positions of all active G proteins. From this a direction estimate is computed by normalizing:

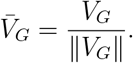

Where appropriate, we will use superscripts 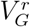 and 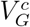 to further distinguish the cases where the vector (B4) is computed under the ratiometric model versus the classical model, respectively. Like the estimates 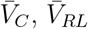 and 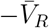, the estimate 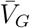 is a random unit vector that depends on time. However, 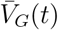 is not a function of the receptor data {*X*_*k*_, *R*_*k*_(*t*)} at any fixed time; 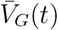 depends on receptor information from previous times, due to the way in which G proteins interact with receptors. We obtain *n* approximately independent estimates for 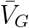 by computing 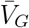 at sufficiently spaced time points *t*_1_ < *t*_2_ < · · · < *t*_*n*_ where *t*_*k*+1_ − *t*_*k*_ ≥ 1*s*.

To compare these direction estimates with the true gradient direction *θ*, we compute the angular deviation arccos 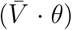 of the estimate 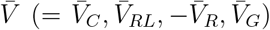 from the true direction *θ*. The empirical cumulative distributions of these angular deviations are shown in (Fig. 1B, 1D, 2D, 3C) of the main text – this function of *α* ∈ [0, 180] is the fraction of the random estimates 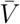 within *α* degrees of the true direction *θ*. In (Fig. 2C, 3A, 3B) of the main text, we also show the fluctuation of these estimates over time, while the receptor and G protein states change asynchronously (but receptor positions are fixed).

### 1. Cramér-Rao bound

The estimates 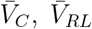 and 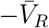 are random variables; in fact, they are functions of the receptor data 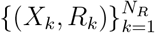 at a fixed time. As such, we examine their efficiency by comparing their variance to theoretical lower bounds provided by the Cramér-Rao theory. The present setting is complicated by the fact that the parameter space for the direction parameter *θ* is the unit sphere 𝒮_1_, which is not a linear space. Nevertheless, the variance of an estimator on the sphere can be bounded from below in terms of the inverse ℐ_*R*_^−1^ of the Fisher information matrix

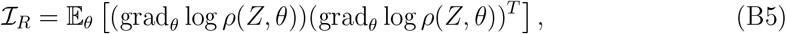

where *ρ*(*Z, θ*) is given by (B1) with 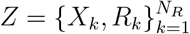, and 𝔼 _*θ*_ denotes integration with respect to density *ρ*(·, *θ*), parameterized by *θ*. This matrix ℐ should be regarded as a symmetric 2 × 2 matrix acting on the tangent space at *θ* ∈ 𝒮_1_. An applicable Cramér-Rao type bound is given in [37]. (See also Chapter 13 of [50]). It follows from Theorem 3.2 and Example 4.2 of [37] that the estimate 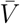 must satisfy

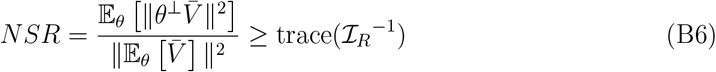

where *θ*^⊥^*Y* = (*I* − *θθ*^*T*^)*Y* is the projection of vector *Y* ∈ ℝ^3^onto the subspace orthogonal to the vector *θ*. By definition 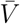is normalized so that 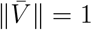; however, 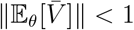. The law of large numbers implies that as 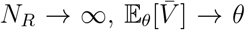 and 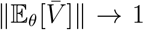. For a weak signal and with few receptors, however, 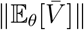 may be small. Nevertheless, (B6) holds as long as the estimator 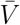 is unbiased and 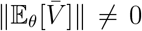, which is the case here for each of the estimators 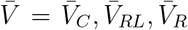, by the symmetry of our choice of *C*(*x, θ*). In particular, 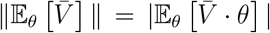 holds in our setting. In the ratio appearing in (B6), 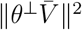 is zero if and only if 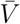 lies in the line spanned by the true direction *θ*; on the other hand, 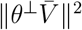 is maximal when 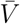 is orthogonal to *θ*. We also note that 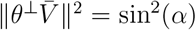 where *α* = arccos 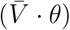 is the angular deviation between 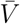 and *θ*.

We refer to the ratio in (B6) as the “noise to signal” ratio (NSR) for the estimate *V*. It is similar to a squared coefficient of variation (*CV*)^2^ for the normalized vector 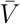: the numerator is the variance of the normalized vector 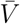 in the directions orthogonal to *θ*, while the denominator is the mean of 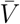 squared. On the other hand, this ratio differs from the usual notion of coefficient of variation in that the quantity 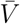 is vector-valued, and the denominator 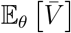 is not the mean of 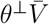. Another Cramér-Rao type bound is given by [51] (see Theorem 2 and Corollary 2):

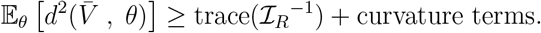

Here *d*(*u, v*) = arccos(*u* · *v*) is the standard Riemannian distance between points *u* and *v* on the unit sphere, and the curvature terms (due to the positive curvature of 𝒮_1_) are negligible when 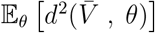 is small (when *N*_*R*_ is sufficiently large).

The Fisher information matrix can be computed, using the fact that the joint density *ρ* in (B1) has a product structure (the pairs {(*X*_*k*_, *R*_*k*_)} are independent and identically distributed). One finds that

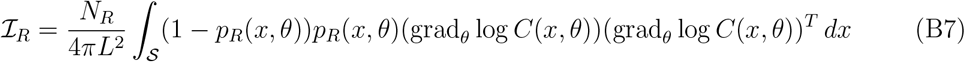

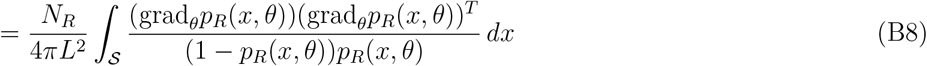

Assuming *p*_*R*_ is a function of *x* · *θ*, then this symmetry of *p*_*R*_ about *θ* implies that ℐ_*R*_ is a multiple of the 2×2 identity matrix. For various choices of *C*(*x, θ*), we evaluate the integral in (B8) numerically, and then compute trace (ℐ_*R*_^−1^) appearing in (B6). For 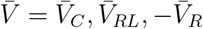, we approximate the ratio

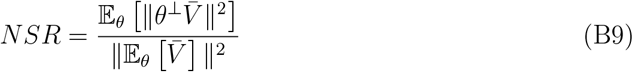

and compare with the lower bound (B6). The expectations are approximated by generating independent samples of 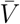 (as described above) and averaging over these samples. The comparisons are shown in (Fig. 1C) of the main text for different total receptor numbers *N*_*R*_.

The G protein configuration at time *t* is described by pairs 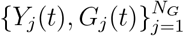. These variables and the corresponding direction estimate 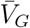 are not a function of the receptor data at any fixed time: they depend on the history of the receptors up to time *t*. Nevertheless, we may ask how the variance of the direction estimate 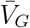 compares to the estimates 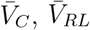 and 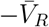, based on the receptor data at a fixed time. To make this comparison, we compute the ratio

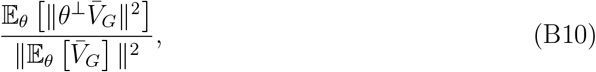

approximating the expectation by averaging over approximately independent samples of 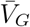. This ratio is shown in (Fig. 2E, 3D, 4A, 4E) of the main text along with the same statistic for the direction estimates 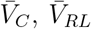 and 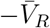. We see that when *N*_*R*_ is small (*N*_*R*_ = 375) and when using the ratiometric model for G protein deactivation, the ratio (B10) is below the lower bound trace (ℐ_*R*_^−1^) in (B6), below the theoretical lower bound for the estimates 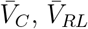, and 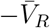 which use receptor data at a fixed time only. This is because 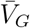 is more concentrated near the true direction *θ* than are the other estimates, even more concentrated than what the theoretical limit allows for any estimator based only on receptor data at a fixed time. The estimator 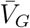 is not a function of the receptor data at any fixed time, so the theoretical bound in (B6) does not apply to the variance ratio (B10) for 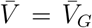. We propose that the fact that the active G protein estimator violates this bound indicates that the G protein configuration is effectively averaging receptor data over a period of time, incorporating an effectively larger amount of receptor information than what is available at a fixed time.

#### 2. Weak signal behavior of Fisher information

Let us suppose *C*(*x, θ*) is given by (A3) with *b* << 1 small. Then grad_*θ*_ log 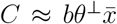 where 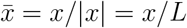. In this case, the Fisher information ℐ_*R*_ is close to a multiple of the 2*x*2 identity matrix, ℐ_*R*_ ≈ *N*_*R*_*κ*^2^*I*, with

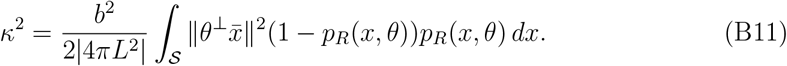

When *b* is small, then *p*_*R*_(*y, θ*) ≈ *C*_0_*/*(*C*_0_ + *K*_*d*_). Hence,

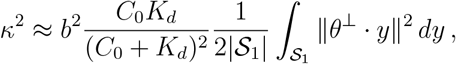

where 𝒮_1_ is the unit sphere. By symmetry, we have 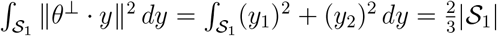. Thus, we have

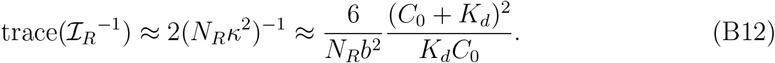

This has the same form as the analogous expression in the two-dimensional model of [32] (c.f. equation (6) for 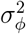 therein, with *p* = 2*b*). In particular, as observed there, this quantity is minimized when *C*_0_ = *K*_*d*_, suggesting that sensing is most effective when *C*_0_ = *K*_*d*_.

Notice that the expression in (B12) diverges as *N*_*R*_*b*^2^ → 0, while the the numerator in (B6) always satisfies 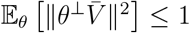. Thus, it must also be true that the denominator in (B6) satisfies 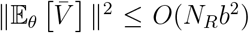 as *b* → 0. (This can be justified directly, by approximating the distribution of *V* (unnormalized) by a Gaussian with mean of *bθ* and covariance 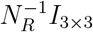.) On the other hand, as 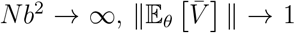. So, (B12) and (B6) together imply that when *b* is small and *N*_*R*_*b*^2^ → ∞,

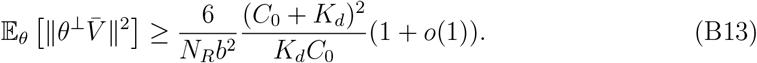

In particular, the mean squared angular deviation of 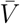is of the order 1*/*(*N*_*R*_*b*^2^). Nevertheless, the bound (B6) holds even when *N*_*R*_*b*^2^ is not large.

#### 3. Covariance and reliability thresholds

How large must *N*_*R*_ be in order that *V*_*RL*_, −*V*_*R*_, or *V*_*C*_ be a reliable estimate of the direction *θ*? The Cramér-Rao bound shows that the mean squared angular deviation of these estimates must be at least of order 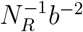. Here we consider an upper bound on angular deviation of these estimates, and we derive an explicit threshold on *N*_*R*_ such that when *N*_*R*_ is above this threshold, the median angle deviation of these estimates will be within a certain tolerance. As we will show, this framework also extends the estimate *V*_*G*_ based on G protein, and it is a useful perspective for comparing *V*_*RL*_, −*V*_*R*_, and *V*_*C*_ with *V*_*G*_. Given the true direction *θ* ∈ *S* of the ligand gradient, and given a tolerance *δ* > 0, we say that a vector *V* (either *V* = *V*_*RL*_, −*V*_*R*_, or *V*_*C*_) is a *δ***-reliable** direction estimate for *θ* if

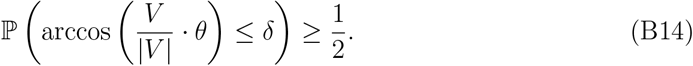

Note that arccos 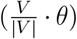 is the angle between the vector *θ* and the vector *V*, even if *V* is not a unit vector. So, condition (B14) is equivalent to saying that the median angle deviation of *V* from *θ* is less than *δ*. Define the cone of angle *δ* around *θ*:

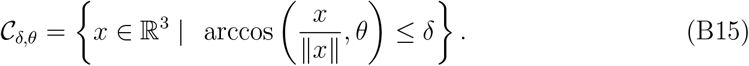

Since the cone 𝒞_*δ,θ*_ is invariant under scaling *x* ↦ *rx* for any *r* > 0, *V* ∈ 𝒞_*δ,θ*_ if and only if 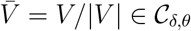. Thus, the reliability condition (B14) is equivalent to

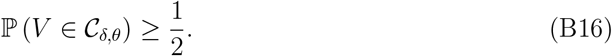

For a fixed deviation tolerance (e.g. *δ* = 30°), how large must *N*_*R*_ be in order that the estimates *V*_*RL*_, −*V*_*R*_, or *V*_*C*_ be reliable in this sense? While the exact distributions of *V*_*RL*_, −*V*_*R*_, and *V*_*C*_ are not explicit, we can estimate ℙ (*V* ∈ 𝒞_*δ,θ*_) in terms of the means and covariances of *V* = *V*_*RL*_, −*V*_*R*_, *V*_*C*_; this will provide a sufficient condition for *δ*-reliability. We say that a random vector *V* is unbiased if its mean *µ* = 𝔼 [*V*] is aligned with *θ*, which means that *µ* = |*µ*|*θ*.

##### Proposition B.1

*Suppose that a random vector V has mean µ* = 𝔼 [*V*] *and covariance matrix* Σ = *Cov*(*V*). *Suppose that V is unbiased: µ* = |*µ*|*θ. If*

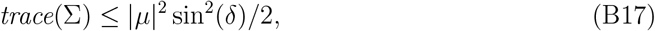

*then V is δ-reliable estimate of θ*.

**Proof:** If *r* ≤ |*µ*| sin(*δ*), then the ball of radius *r* centered at *µ* is contained in the cone *C*_*δ,θ*_. See Figure 8. In other words, the condition

**FIG. 8.**
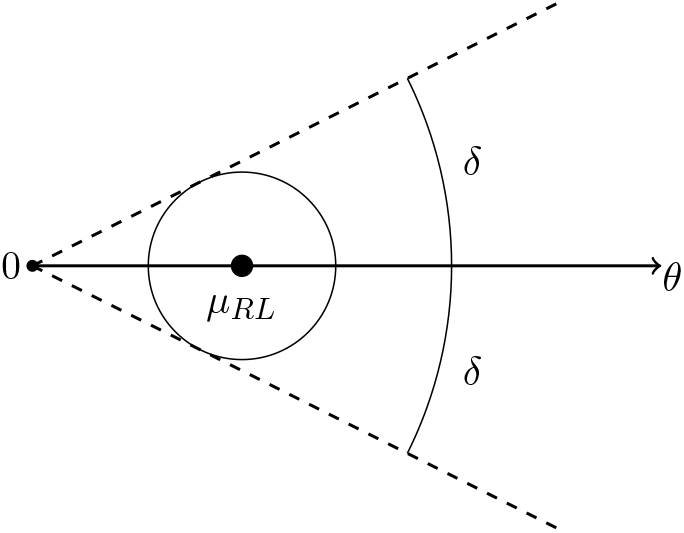
The figure illustrates the cone *C*_*δ,θ*_. The ball centered at *µ* has radius *r* = |*µ*| sin(*δ*). The condition (B17) guarantees that the standard deviation of *V* is less than 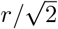, so that *V* lies in the ball with probability at least 1*/*2.

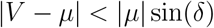

implies *V* ∈ 𝒞_*δ,θ*_. By Chebyshev’s inequality:

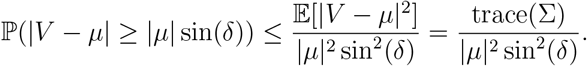

If (B17) holds, then the right side is less than 1*/*2. Hence

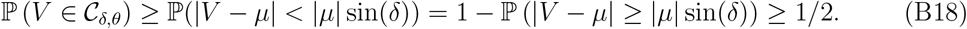

Thus, *V* is *δ*-reliable.

The condition (B17) is a sufficient condition for *V* to be *δ*-reliable, but it is not a necessary condition. Nevertheless, condition (B17) is easier to verify than computing the probability in (B16), and it is not based on any approximation.

We now apply this to the particular cases *V* = *V*_*RL*_, −*V*_*R*_, *V*_*C*_. Let *µ*_*RL*_ denote the mean of *V*_*RL*_:

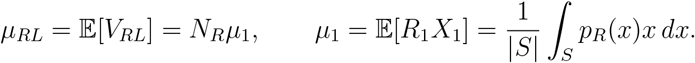

The mean of −*V*_*R*_ is the same, since 𝔼 [*X*_*i*_] = 0:

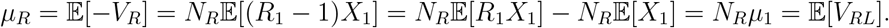

Since the pairs 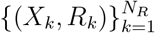 are independent, the covariance matrix of *V*_*RL*_ is

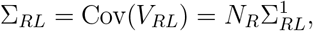

where 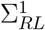 is the covariance of *R*_1_*X*_1_:

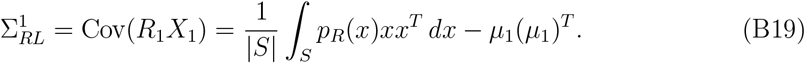

Similarly, the covariance matrix of −*V*_*R*_ is

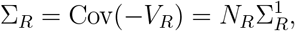

where 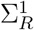 is the covariance of (*R*_1_ − 1)*X*_1_:

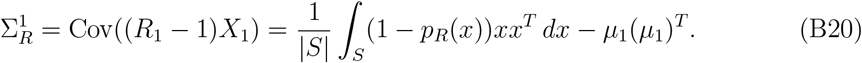

The mean of *µ*_*C*_ of *V*_*C*_ is exactly twice the mean of *V*_*RL*_:

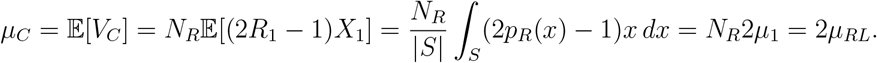

The covariance of *V*_*C*_ is

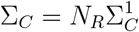

where 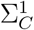 is the covariance of (2*R* − 1)*X*_1_:

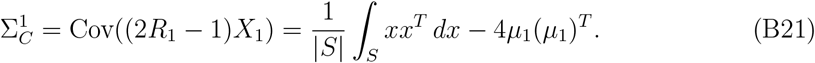

Notice that the relation

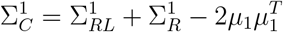

always holds. The estimates *V*_*RL*_, −*V*_*R*_, and *V*_*C*_ are all unbiased if *µ*_1_ is aligned with *θ*: *µ*_1_ = |*µ*_1_|*θ*.

By applying Proposition B.1 with *V* = *V*_*RL*_, −*V*_*R*_, *V*_*C*_ we obtain the following thresholds for *δ*-reliability:

##### Corollary B.1

*Suppose that the estimates V*_*RL*_, −*V*_*R*_, *and V*_*C*_ *are unbiased: µ*_1_ = |*µ*_1_|*θ. The reliability condition* (B17) *for V*_*RL*_ *is*

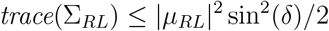

*or, equivalently*,

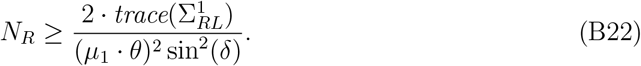

*The reliability condition* (B17) *for* −*V*_*R*_ *is*

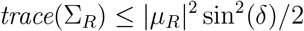

*or, equivalently*,

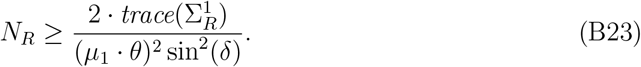

*The reliability condition* (B17) *for V*_*C*_ *is*

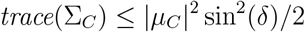

*or, equivalently*,

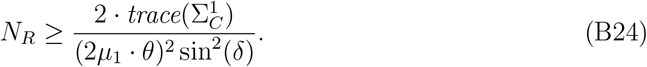

We may interpret the quantity *µ*_1_ · *θ* to be the “signal strength”; this calculation shows that reliability of *V*_*RL*_, −*V*_*R*_ and *V*_*C*_ requires that *N*_*R*_ is of the order (*µ*_1_ · *θ*)^−2^, which is large when the signal is weak (shallow ligand gradient). Suppose that the gradient is weak: *C*(*x*) = *C*_0_(1 + *bx* · *θ*) with *b* > 0 small (here we assume *L* = 1). Then *p*_*R*_(*x*) is close to linear in *x*:

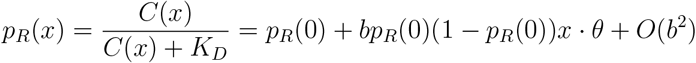

and thus

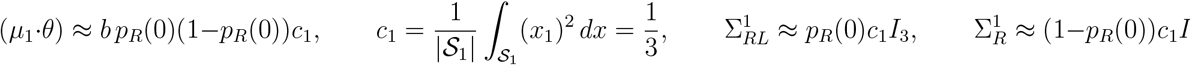

where *I*_3_ is the 3 × 3 identity matrix. Hence, in this weak gradient regime, the sufficient condition (B22) for *V*_*RL*_ to be *δ*-reliable reduces to

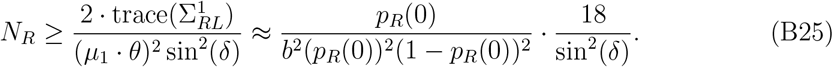

The condition (B23) for −*V*_*R*_ to be *δ*-reliable reduces to

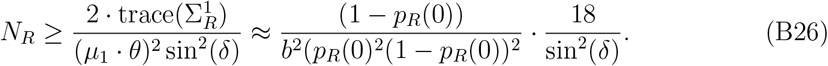

The condition (B24) for *V*_*C*_ to be *δ*-reliable reduces to

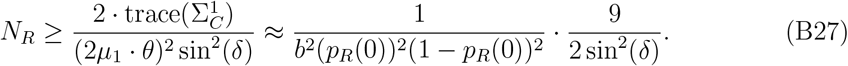

Comparing (B25), (B26), and (B27), we see that when *p*_*R*_(0) ∈ (1*/*4, 3*/*4), *V*_*C*_ has the lowest threshold for reliability. When *p*_*R*_(0) = 1*/*2 (i.e. *C*_0_ = *K*_*d*_), the thresholds (B25) and (B26) for *V*_*RL*_ and −*V*_*R*_ are the same, and the threshold (B27) for *V*_*C*_ is exactly ½ of (B25) or (B26). In this scenario, *V*_*C*_ is effectively using twice the data as *V*_*RL*_ or −*V*_*R*_. For *p*_*R*_(0) outside of the range (1*/*4, 3*/*4), *V*_*C*_ is not the most reliable of the three estimates. When *p*_*R*_(0) < 1*/*4 (i.e. *C*_0_ < (1*/*3)*K*_*d*_), the reliability threshold for *V*_*RL*_ ((B25)) is lowest among the three; when *p*_*R*_(0) > 3*/*4 (i.e. *C*_0_ > 3*K*_*d*_), the reliability threshold for −*V*_*R*_ ((B25)) is lowest among the three. The intuition about why *V*_*C*_ would perform worse than *V*_*RL*_ when *p*_*R*_(0) < 1*/*4 is that in this regime the majority of receptors are inactive and their conditional angular distribution is more uniform than the conditional distribution of active receptor. Indeed, the conditional density of a receptor’s location, conditioned on it being inactive, is proportional to 1 − *p*_*R*_(*x*) = *K*_*d*_*/*(*K*_*d*_ + *C*(*x*)), and this density is more uniform when *C*_0_*/K*_*d*_ is small. So, when *C*_0_*/K*_*d*_ is small, the estimate *V*_*C*_ is dominated by the contribution from inactive receptors which are approximately uniformly distributed on the sphere.

Another approach to estimating the probability in (B16) would be to approximate *V* by a Gaussian random vector having the same mean and covariance (justified by the central limit theorem when *N*_R_ is large). If *V* has mean |*µ*|*θ* and covariance matrix 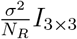, then under this approximation

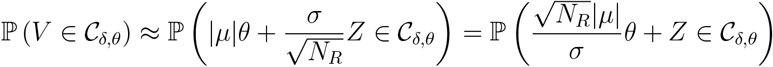

where *Z* is a standard Gaussian vector in ℝ^3^ with mean zero and covariance *I*_3×3_. Thus, *V* is *δ*-reliable once 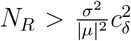, where the constant *c*_*δ*_ > 0 is the smallest value *r* such that ℙ (*rθ* + *Z* ∈ 𝒞_*δ,θ*_) ≥ 1*/*2. Applied to *V*_*RL*_, −*V*_*R*_, and *V*_*C*_, this approach yields thresholds with the same scaling as (B25), (B26), (B27), but with the less explicit universal constant 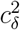.

## Appendix C: Time averaging

In the ratiometric model, the instantaneous G protein states (active/inactive) are derived from historical receptor states. Each G protein’s state is determined by its most recent interaction with a receptor, either active or inactive (Figure 9). If *N*_*G*_ > *N*_*R*_, this mechanism may allow the G protein states to represent more information than what is encoded by the receptor states at a single time. Here we explain this mechanism. Throughout this section we consider only the ratiometric model for G protein dynamics.

**FIG. 9.**
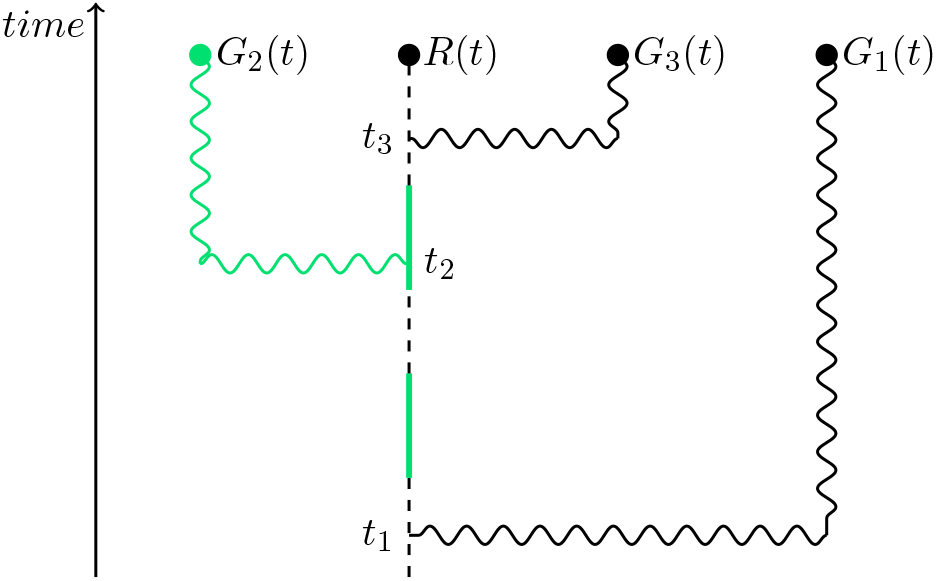
This figure illustrates how G proteins record the historical state of a receptor. The states of G proteins *G*_1_, *G*_2_, and *G*_3_ at time *t* were obtained through interaction with the same receptor, but at different earlier times. The protein *G*_1_ records the receptor state from time *t*_1_ < *t*; we say at that time *t* the age of *G*_1_ is *t* − *t*_1_. Protein *G*_2_ records the receptor state from time *t*_2_ < *t*; at time *t* the age of *G*_2_ is *t* − *t*_2_. Even though each protein last interacted with the same receptor, the states they record at time *t* may differ: *G*_1_ and *G*_3_ are inactive, while *G*_2_ is active.

### 1. Sampling the state of a single receptor at multiple random times

For simplicity, we first consider the case of a single receptor, and *n* G proteins interacting with it, and we ignore the spatial aspect of the model for now. The state of a given G protein is determined by the state of the receptor at the time of the protein’s most recent encounter with that receptor. If the times at which the *n* proteins most recently interacted with that receptor are *t*_1_, *t*_2_, …, *t*_*n*_ then the states of the G proteins represent *n* samples of the historical state of the (single) receptor. These historical states are correlated, however; the correlation is higher when the interaction times are close together. If the interaction times are far apart, then sampling the state of the receptor at *n* different times can be similar to sampling *n* independent receptors.

The receptor state is a two-state, continuous-time Markov process *R*(*t*); the two states are the active state (1) and the inactive state (0). The transition rates for this chain are given by (A1), where *x* ∈ 𝒮 is the fixed location of the receptor. For clarity in notation we write λ_+_ = *k*_on_*C*(*x*) for the activation rate, λ_−_ = *k*_off_ for the inactivation rate, and we define λ = λ_+_ + λ_−_. The stationary distribution for this process is ν = (ν(0), ν(1)), with

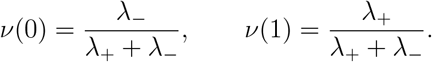

So, ν(0) and ν(1) are the steady-state probabilities of finding the receptor in states 0 or 1, respectively. Let *p*_*t*_(*z, y*) denote the transition probabilities: *p*_*t*_(*z, y*) is the probability that the state at time *t* > 0 is *y*, given that the state at time 0 is *z, y, z* ∈ {0, 1}. These probabilities satisfy the system

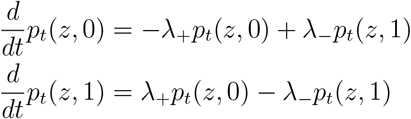

for *x* ∈ {0, 1}. This has an explicit solution. We find that the transition probabilities are

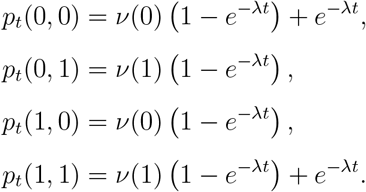

In particular, regardless of the initial state *x, p*_*t*_(*z, y*) converges exponentially fast to ν(*y*), at rate λ = λ_+_ + λ_−_. Thus, λ^−1^ = (*k*_on_*C* + *k*_off_)^−1^ is the time scale for “loss of memory” or mixing of the receptor state.

Now, suppose the receptor process *R* has achieved stationarity, and suppose there are *n* G proteins that read the state of this single receptor at random times *t*_1_, …, *t*_*n*_ (unordered). Assume these times are independent of each other and of the receptor process. Suppose that for each *k, t*_*k*_ is exponentially distributed with mean *τ* = 𝔼 [*t*_*k*_]. We will use *G*_*i*_ ∈ {0, 1} to denote the state of the *i*th G protein (i.e. *G*_*i*_ = *R*(*t*_*i*_)). Although the times 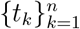 are independent, the random variables (*G*_1_, …, *G*_*n*_) = (*R*(*t*_1_), …, *R*(*t*_*n*_)) are dependent, since the data come from the same receptor. However, the level of dependence is controlled by the dimensionless parameter *λτ*, as we now show. Let *A*_*G*_ denote the fraction of receptor readings that are in the active state:

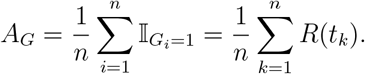

Let 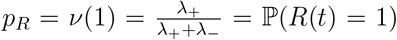 be the probability that the receptor is active (*R* is statistically stationary, so this probability does not change with time). The autocorrelation function for the receptor state is: 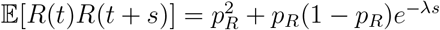.

#### Proposition C.1

*The mean and variance of A*_*G*_ *are* 𝔼 [*A*_*G*_] = *p*_*R*_ *and*

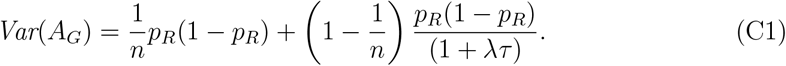

The first term in this expression, *p*_*R*_(1 − *p*_*R*_)*/n*, would be the variance if the *n* readings were independent (e.g. taken separately from *n* independent receptors); in the limit *n* → ∞ this term vanishes. The second term in the variance is the result of temporal correlation in the receptor process. As *λτ* → ∞, the last term vanishes, leaving only *p*_*R*_(1 − *p*_*R*_)*/n*.

#### 2. Effective receptor number

Let us compare this result to an alternative model, which will motivate the notion of **effective receptor number**. Instead of there being a single receptor with state changing in time, let us now suppose there is a cluster of *n*_*R*_ receptors with independent states that do not change in time. Let *p*_*R*_ be the probability that a given receptor in the cluster is active. Suppose that each of the *n* G proteins samples a state from this cluster: each protein chooses one of the *n*_*R*_ receptors at random and records its state. So, two proteins that choose the same receptor must record the same state. As before, let *A*_*G*_ be the fraction of readings that are in the active state:

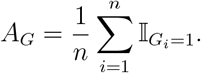

##### Proposition C.2

*The mean and variance of A*_*G*_ *are* 𝔼 [*A*_*G*_] = *p*_*R*_ *and*

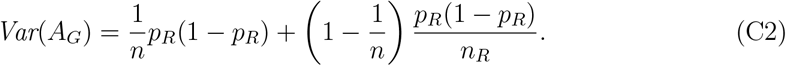

Both models have the same mean *p*_*R*_ for all parameter choices. Comparing (C1) and (C2), we see that the variance in the two models is the same when (1 + *λτ*) = *n*_*R*_. Thus, we could think of (1 + *λτ*) as an **effective number of receptors**: when the G proteins read a single receptor at independent random times, it is as if each G protein reads a state chosen randomly and independently from a group of (1 + *λτ*) independent receptors. The fact that the G proteins read historical states effectively expands the pool of independent receptors. Interestingly, the quantity *n*_*R*_ = (1 + *λτ*) does not depend on *n*, assuming *n* > 1.

#### 3. Derivation of Proposition C.1 and Proposition C.2

For both models, the variance of *A*_*G*_ is

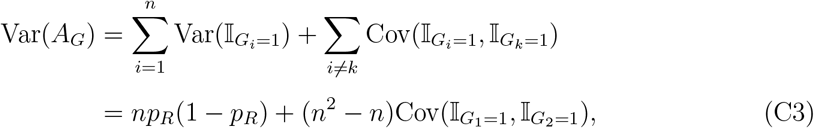

and the covariance term is

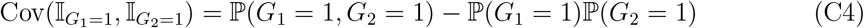

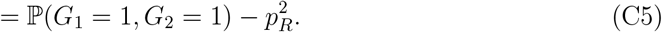

For the model considred in Proposition C.1, the joint density of the independent times (*t*_1_, *t*_2_) is 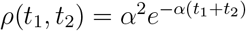 for *t*_1_, *t*_2_ ≥ 0. Therefore,

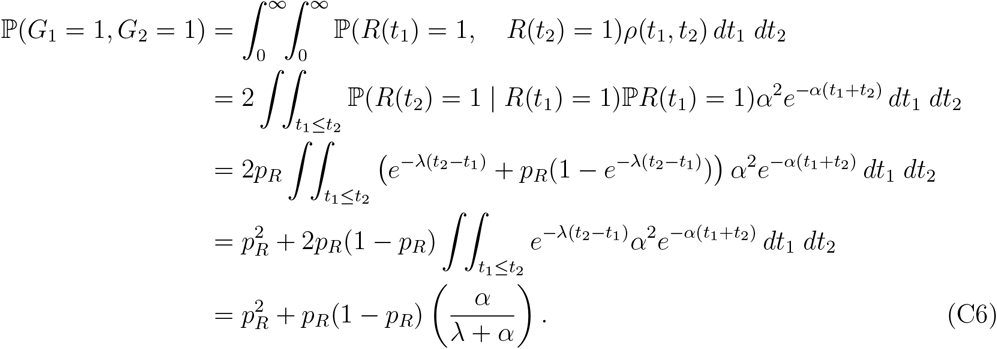

Therefore,

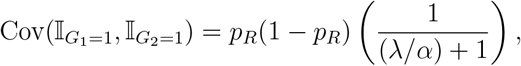

and

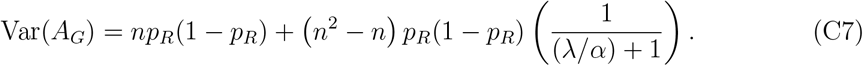

For the model considered in Proposition C.2,

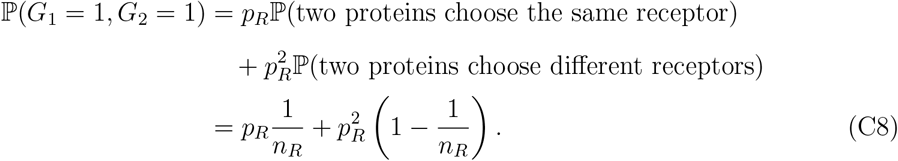

Therefore,

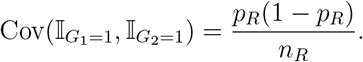

## Appendix D: Spatially extended, ratiometric model

In the analysis above we considered a population of G proteins interacting with a single receptor. Here we show how these ideas extend to the case when there are multiple receptors distributed spatially across the cell surface, and the G proteins interact with them, according to the ratiometric model, as the proteins diffuse over the surface. Throughout this section, we assume the ratiometric model, so 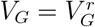.

### 1. G protein ages

Considering the times at which a G protein interacts with a receptor, we define the **age** of a G protein at time *t* to be *s* ≥ 0 if the last interaction between that G protein and some receptor occured at time *t* − *s*. If the *G* protein has age *s*, then that G protein has been in the same state for at least time *t* − *s*. Here we say “at least” time *t* − *s* because it could be that the G protein was already in that state just before this most recent encounter with a receptor. For example, when an already-active G protein encounters an active receptor, the G protein’s age will reset to zero but its state does not change. (See Figure 9). We will say that a G protein at time *t* is **linked** to a particular receptor if that G protein’s last interaction was with that particular receptor – if that G protein’s state at time *t* was recorded from that particular receptor (at time *t* − *s*, where *s* is the protein’s age). Each G protein is linked to only one receptor at a time. Two or more G proteins may be linked to the same receptor, which creates dependence between those G protein states. (See Figure 10). In fact, this linking is the only source of statistical dependence between G protein states: G protein states are dependent only through interaction with the same receptor. Conditioned on being linked to two different receptors, two G protein states are statistically independent. However, for G proteins linked to the same receptor, their ages may be different, and the observations above suggest that if these ages are sufficiently far apart, then the recorded states are approximately independent. As explained above in Proposition C.1, the level of dependence/independence between proteins linked to the same receptor is controlled by the ratio of two times scales – the mean G protein age and the receptor turnover time.

**FIG. 10.**
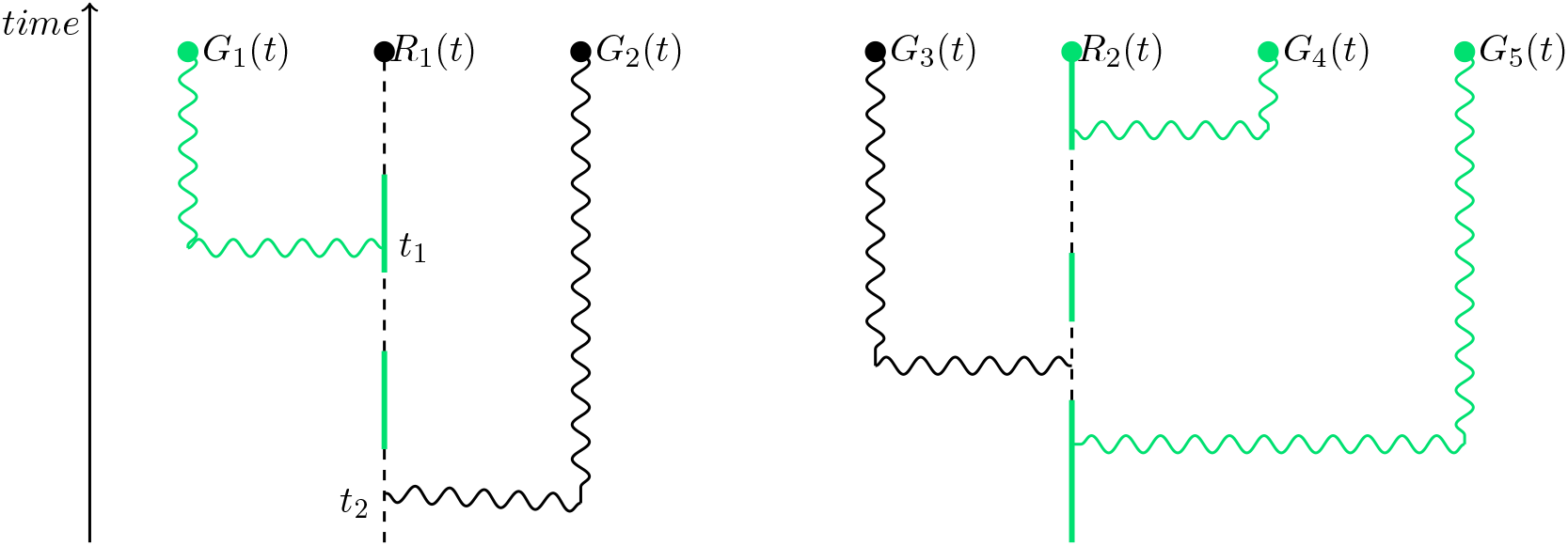
This figure illustrates the most recent receptor-interactions among a collection of five G proteins. The states of G proteins *G*_1_ and *G*_2_ at time *t* were obtained through interaction with the same receptor *R*_1_. Proteins *G*_1_ and *G*_2_ are linked to receptor *R*_1_. However, proteins *G*_3_, *G*_4_, *G*_5_ are linked to receptor *R*_2_, since their most recent receptor interaction was with receptor *R*_2_.

We can estimate the typical age of a G protein. If there are a total of *N*_*R*_ receptors, then the typical distance between receptors is of the order 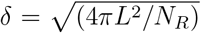 where *L* is the cell radius. This is because 4*πL*^2^*/N*_*R*_ is the surface area per receptor, and the surface is two-dimensional (hence, the square root). Therefore, a G protein diffusing on the cell surface is navigating among receptors that are typically distance *δ* apart. If we want to know when a given G protein last interacted with some receptor (any receptor), it is the same as looking *backwards* in time and asking when does a Brownian motion first hit an *r*_∗_-neighborhood of some receptor, among the *N*_*R*_ receptors distributed across the surface approximately *δ* apart. If the receptors have an interaction radius of *r*_∗_ ≪ *δ* and the G protein diffusion coefficient is *D*_*G*_, then the typical **age** of a G protein will be of the order

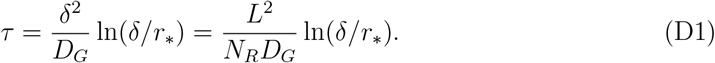

We may also describe this quantity as the time scale of diffusive encounters between a G protein and receptors. This is motivated by thinking about a periodic array of discs in ℝ^2^, having radius *r*_∗_ and with centers *δ* apart. The quantity 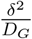 ln(*δ/r*_∗_) is the typical time it takes for a Brownian motion (with diffusion coefficient *D*_*G*_) to hit one of the discs (any disc), when started from a random location. (The logarithm is an artifact of the 2-dimensional nature of the cell surface and of the Green’s function in 2-dimensions.) Here and elsewhere we drop constants of proportionality, such as 4*π*, that do not depend on the parameters, since we are mainly interested in the scaling with parameters. This time *τ* is not the typical time required for a given G protein to find a *particular* receptor. Rather, it is the time required (looking backward in time) to encounter any one of the receptors – this later quantity is what determines the age of a G protein. The relation in (D1) can be derived as in Appendix B of [28]; ignoring constants of proportionality that do not depend on the parameters, (D1) above has the same form as Eq. 14 from [28]. See also [52], [53], and references therein for a broader discussion of first-passage time calculations. This suggests that the age of a G protein is a random variable that is approximately exponentially distributed with mean *τ*. If the number of G proteins is *N*_*G*_ > *N*_*R*_, then on average there are (*N*_*G*_*/N*_*R*_) G proteins per receptor. So, if the receptors and proteins are uniformly distributed on the sphere, we should expect that for each of the *N*_*R*_ receptors there are about (*N*_*G*_*/N*_*R*_) G proteins that are **linked** with that receptor, in the sense defined above.

#### 2. Covariance for G protein, fixed receptor locations

As we did with the simple single-receptor model above, we now consider how the time *τ* effects the variation of the G protein signal. Specifically, we show how for the ratiometric model the covariance of *V*_*G*_ depends on the time scale *τ* and the time scale of receptor dynamics. Here we suppose that the receptor locations are fixed at 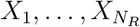. The receptor states are changing randomly, but their locations are fixed. A receptor at *X*_*i*_ activates at rate *k*_on_*C*(*X*_*i*_) and deactivates at rate *k*_off_; thus, the time scale for receptor dynamics at location *X*_*i*_ is λ^−1^(*X*_*i*_) = (*k*_on_*C*(*X*_*i*_) + *k*_off_)^−1^. Suppose there are *N*_*G*_ proteins that are placed at locations 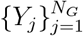, which are independent and uniformly distributed on the sphere. We consider their states 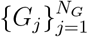 at a fixed time, where *G*_*j*_ ∈ {0, 1}. We are interested in the vector

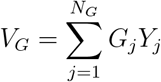

which is the sum of the positions of active G proteins. Let *µ*_*G*_ = 𝔼 [*V*_*G*_] be the mean of *V*_*G*_, and let Σ_*G*_ = Cov(*V*_*G*_) be the covariance matrix. As in (B16), we say that the G protein signal *V*_*G*_ is a *δ***-reliable** estimate if

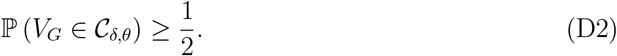

Proposition B.1 applied with *V* = *V*_*G*_ gives us a sufficient condition for *δ*-reliability of *V*_*G*_:

##### Corollary D.1

*Suppose that the estimate V*_*G*_ *is unbiased. If trace*(Σ_*G*_) ≤ |*µ*_*G*_|^2^ sin^2^(*δ*)*/*2, *then V*_*G*_ *is δ-reliable*.

To make use of this criteria and to compare the reliability of *V*_*G*_ with the reliability of *V*_*RL*_, we need to compute *µ*_*G*_ and Σ_*G*_. We now explain how these can be calculated for the ratiometric model.

The state *G*_*j*_ of the protein at position *Y*_*j*_ was read from a particular receptor at some previous time – we say that this protein is **linked** to that particular receptor. Define random variables 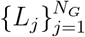 which indicate the receptor to which the *j*th protein is linked: *L*_*j*_ = *k* means that the *j*th protein (at *Y*_*j*_) is linked to the *k*th receptor (which is at position *X*_*k*_). Possibly multiple G proteins can be linked to the same receptor, but they may or may not read the same state from that receptor (because they interacted with that receptor at different times, and the receptor state is changing over time).

Let *η*_*i*_(*y*) denote the conditional density of *Y*_*j*_, given *L*_*j*_ = *i*. This *η*_*i*_ is depends on the geometric arrangement of all receptors 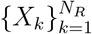, and we denote this dependence by *η*_*i*_([*X*], *y*). Although this quantity is impossible to compute exactly/explicitly, we should think of this as being concentrated in a neighborhood of *X*_*i*_, as G proteins are most likely to be linked to nearby receptors. Thus,

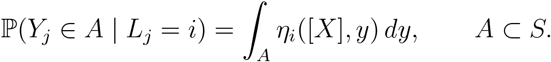

Given *L*_*j*_ = *i*, the *j*th G protein takes its state from the *i*th receptor at some time *T*_*j*_. Motivated by the analysis above, we make the simplifying assumption that this time (the age of that protein) is exponentially distributed with mean *τ, T*_*j*_ ~ Exponential(1*/τ*), where the parameter *τ* is given by (D1). We also assume that the ages are independent of G protein positions.

##### Proposition D.1

*Assume the ratiometric model for G protein dynamics. Given fixed receptor locations* 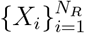, *the mean of V*_*G*_ *is*

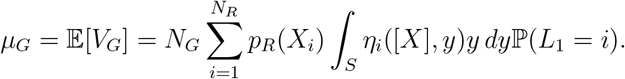

*The covariance matrix* Σ_*G*_ = 𝔼 [(*V*_*G*_ − (*V*_*G*_))(*V*_*G*_ − (*V*_*G*_))^*T*^] *of V*_*G*_ *is*

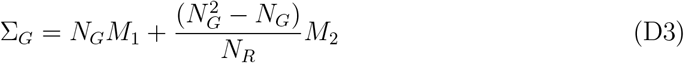

*Where*

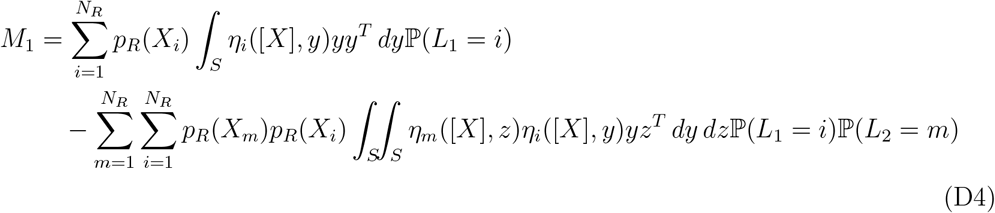

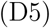

*and*

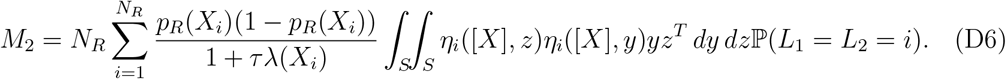

The quantity

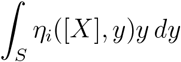

in the expression for *µ*_*G*_ is the mean location of a G protein linked to the *i*^*th*^ receptor (at location *X*_*i*_). Observe that the covariance matrix Σ_*G*_ is the sum of two terms. The second term, *M*_2_, results from correlations caused by G proteins possibly interacting with the same receptor. This *M*_2_ is the term where time-scales play a role, through *τλ*(*X*_*i*_). When *τλ* is large, meaning that receptor dynamics are fast relative to G protein ages, then *M*_2_ may be negligible.

From Proposition D.1 we can identify two parameter regimes corresponding to different behavior of the random fluctuation of *V*_*G*_. The covariance of 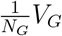 is

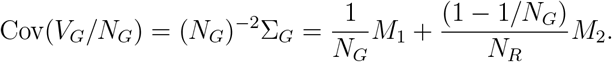

Recall that the matrix *M*_2_ depends on *τ* and λ. The quantity λ(*x*) = *k*_on_*C*(*x*) + *k*_off_ depends on *x*. Nevertheless, when the gradient is shallow, we have λ ≈ λ_0_ = *k*_on_*C*_0_ + *k*_off_. Thus, Proposition D.1 shows that the dimensionless parameter λ_0_*τ* = (*k*_on_*C*_0_ +*k*_off_)*τ*, which is ratio of the G protein age *τ* with the receptor turnover time 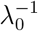, plays an important role. (The product λ_0_*τ* is analogous to a Damköhler number.) This analysis suggests two qualitatively different regimes for the behavior of the covariance of *V*_*G*_:

- **The** *G***-limited regime**. If

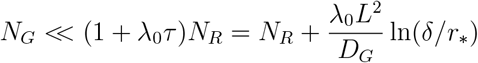

then we have

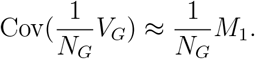

In this regime, the covariance is approximately that of *N*_*G*_ proteins that are sampling independent receptors. Either *N*_*R*_ is large or λ_0_*τ* is large, so that the receptor states sampled by the G proteins are approximately independent. Consequently, increasing *N*_*R*_ or λ_0_*τ* further does not decrease the covariance as significantly as would an increase in *N*_*G*_. In other words, small *N*_*G*_ is the main limitation for reliability of *V*_*G*_ in this regime. Note: it is possible that *N*_*R*_ < *N*_*G*_ < (1 + λ_0_*τ*)*N*_*R*_. In other words, even if *N*_*R*_ < *N*_*G*_, the time scales may be such that the covariance of *V*_*G*_ is relatively insensitive to changes in *N*_*R*_ because (1 + λ_0_*τ*)*N*_*R*_ >> *N*_*G*_. This is consistent with our observation that the estimate 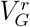 may be robust even at low *N*_*R*_, as illustrated in Figure 2 and 3 of the main text. In particular, in this regime the noise-to-signal ratio for 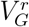 may be smaller than trace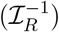 from (B6), which is the theoretical lower bound that applies to *V*_*C*_ and to any unbiased estimate based on instantaneous receptor states at a single fixed time.
- **The Receptor-limited regime**. If

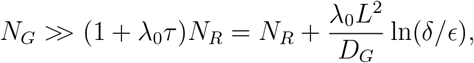

then we have

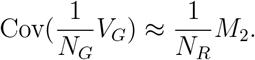

In this regime, there are typically many G proteins linked to a given receptor. Variation of the estimate *V*_*G*_ is dominated by fluctuation in the receptor states (which are passed on to the G protein), and there is a relatively high level of correlation in the receptor states recorded by the G proteins linked to the same receptor. Increasing *N*_*G*_ further does not decrease the covariance as significantly as would an increase in either *N*_*R*_ or λ_0_*τ*. In other words, even though *N*_*G*_ is large, *N*_*R*_ is the main limitation for reliability of *V*_*G*_ in this regime.

In view of these calculations, we define the quantity

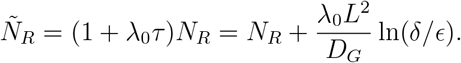

We call this the **effective receptor number** – this number represents an effective number of receptor states made available through historical sampling. The quantity *Ñ*_*R*_ should not be thought of as something that applies to a particular cell. Instead, *Ñ*_*R*_ can be regarded as a property of the ratiometric model, a model whose statistical properties are completely determined by parameters *N*_*R*_, *N*_*G*_, *D*_*G*_, *L, r*_∗_, *k*_off_, *K*_*d*_ and the ligand concentration *C*(*x*).

Considering how these two regimes depend on the parameters, we make the following observations:

- Regarding *D*_*G*_: Faster diffusion of G protein (*D*_*G*_ large) reduces *τ*. This is because when the diffusion of G protein is fast, the proteins typically interact with receptors very frequently, and their age is lower, so their states are more correlated with other G proteins that also hit the same receptor in the very recent past (the states of linked G proteins are likely to coincide). In the hypothetical extreme *D*_*G*_ → ∞, the age of a G protein goes to zero so that the G proteins are recording only information from the present state of the receptors. In this case, *Ñ*_*R*_ ≈ *N*_*R*_. Conversely, slower diffusion (*D*_*G*_ small) increases *τ* and the effective receptor number *Ñ*_*R*_. This is because with slow diffusion of G protein, their ages tend to be larger and further apart, so even the states of linked G proteins (i.e. G proteins that are linked to the same receptor) are approximately independent.
- Regarding λ_0_: Faster receptor dynamics λ_0_ >> 1 increases *Ñ*_*R*_. This is because the states of a given receptor *R*(*t*_1_), …, *R*(*t*_*n*_) at some times *t*_1_ < *t*_2_ < · · · < *t*_*n*_ (that do not depend on λ_0_) will be approximately independent when λ_0_ is large. Thus, if multiple G proteins obtained their state by last interacting with that same receptor, then those proteins are encoding approximately independent state values. Conversely, slow receptor dynamics (λ_0_ << 1) decreases *Ñ*_*R*_. A key point here is that changing the receptor turnover time 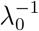 has no effect on the distribution of G protein ages.
- Regarding *r*_∗_: decreasing the interaction radius *r*_∗_ will tend to increase G protein ages relative to the receptor turnover time, since it is more difficult for a protein to reach a receptor. This leads to greater independence in the states encoded by the G protein ensemble.

Thus far we have reasoned under the assumption that ligand concentration *C*(*x*) is not far from uniform:

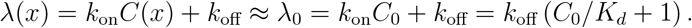

With non-uniform ligand concentration, the receptor turnover time depends on position. So, in principle, the G protein on front end of the cell (where ligand concentration *C*(*θ*) is highest) should encode a higher level of independence than on the back end, because λ is larger on the front end. However, in the scenarios we are considering the ratio (*C*(*θ*) + *K*_*d*_)*/*(*C*(−*θ*) + *K*_*d*_)), with *θ* denoting the gradient direction, is not very large. So, the difference in this effect between front and back may not be significant.

#### 3. Comparing 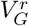 with *V*_*RL*_ when *N*_*R*_ is sufficiently large

These expressions for the mean *µ*_*G*_ and covariance Σ_*G*_ of 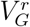 are complicated. In particular, explicitly computing the probability of linking to a particular receptor is not feasible, as it depends on the geometric configuration of all receptors. Nevertheless, we can gain further insight by making a simplifying assumption that *N*_*R*_ is large and the receptors are evenly distributed over *S*. This will allow us to simplify *µ*_*G*_ and Σ_*G*_ and compare them with *µ*_*R*_ and Σ_*R*_, the mean and covariance of *V*_*RL*_. In particular, we will show that in certain parameter regimes, 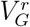 may be a more reliable estimate of *θ* than *V*_*RL*_. This analysis is consistent with the results of our numerical simulations.

Under the assumption that *N*_*R*_ is large and the receptor locations are well-spaced over *S*, we expect that a G protein placed uniformly at random on *S* is equally likely to link with any receptor:

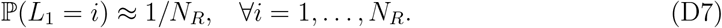

Furthermore, under this assumption we expect that the conditional density *y* ↦ *η*_*i*_([*X*], *y*) is concentrated in a small neighborhood of *X*_*i*_. Thus, we approximate

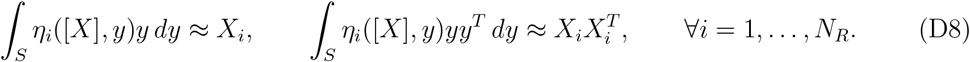

Using the approximations (D7) and (D8) we have the following approximations of the mean and covariance of 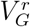. Recall from (B20), that *V*_*RL*_ satisfies

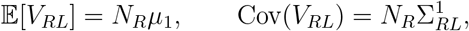

Where

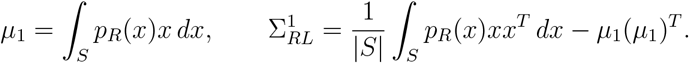

##### Proposition D.2

*Using the approximations* (D7) *and* (D8) *and assuming N*_*R*_ *is large, the mean µ*_*G*_ *and covariance* Σ_*G*_ *of* 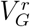 *are*

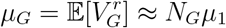

*And*

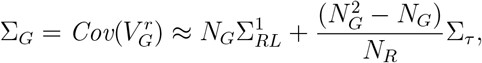

*where*

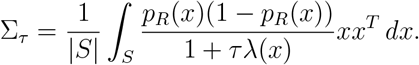

Using this, we can derive the following remarkable consequence about reliability of 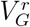 in the G-limited regime:

**Corollary D.2** *When N*_*G*_ > *N*_*R*_ *and τλ is large, the estimate* 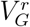 *of θ may be δ-reliable, even while V*_*RL*_ *is not δ-reliable*.

**Proof:** Using these approximations for *µ*_*G*_ and Σ_*G*_, the condition given in Corollary D.1 for *δ*-reliability of 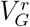 is (approximately):

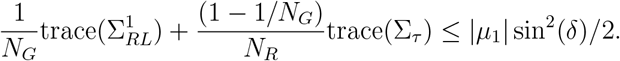

In particular, this holds if

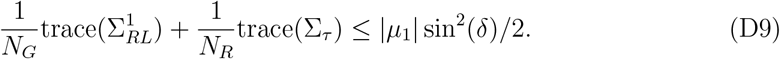

On the other hand, the *δ*-reliability condition for *V*_*RL*_, from Corollary B.1 is

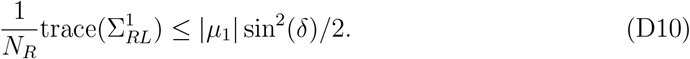

The matrix Σ_*τ*_ depends on *τλ*, but 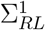 does not. For any *N*_*R*_, we may take *δ* small enough so that *V*_*RL*_ is **not** *δ*-reliable. In particular, (D10) does not hold if *δ* is small. For this same value of *N*_*R*_ and *δ*, however, we may take *N*_*G*_ > *N*_*R*_, and then *λτ* large enough so that the inequality (D9) holds, implying that 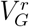 is *δ*-reliable, even though *V*_*RL*_ is not. In particular, it would be necessary for trace(Σ_*τ*_) ≤ (1 − *N*_*R*_*/N*_*G*_)trace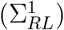.

#### 4. Derivation of Proposition D.1

The mean of *V*_*G*_ is:

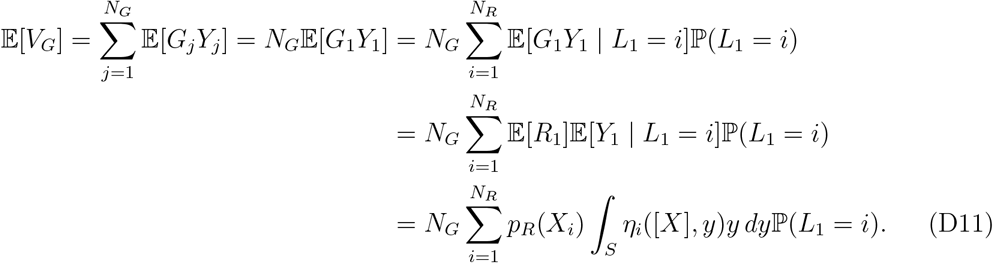

The covariance matrix Σ_*G*_ = 𝔼 [(*V*_*G*_ − (*V*_*G*_))(*V*_*G*_ − (*V*_*G*_))^*T*^] is:

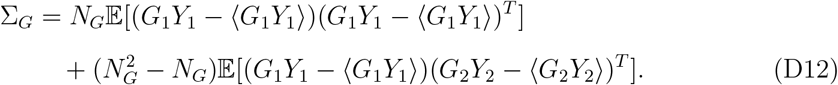

The first term is:

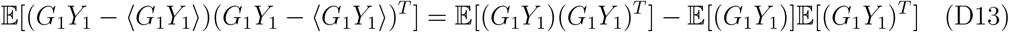

and

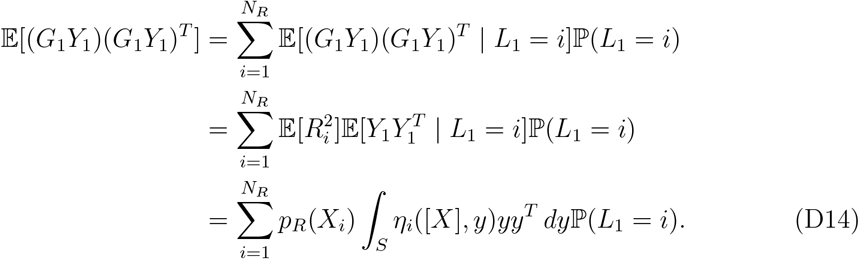

Next,

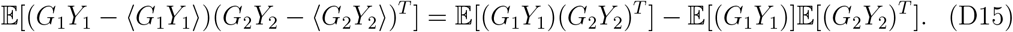

and

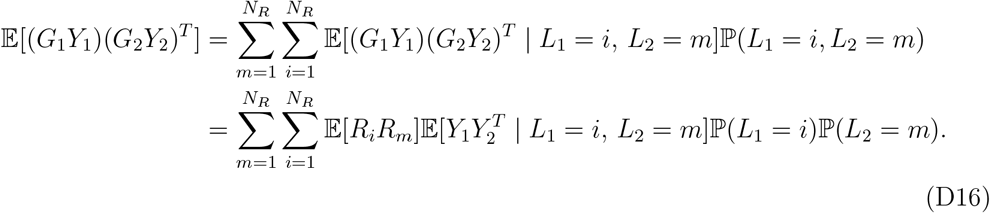

The case *i* ≠ *m* means that the two G proteins are linked to a different receptor (whose states are independent). In this case, 𝔼 [*R*_*i*_*R*_*m*_] = 𝔼 [*R*_*i*_] 𝔼 [*R*_*m*_] = *p*_*R*_(*X*_*i*_)*p*_*R*_(*X*_*m*_). However, the case *i* = *m* means that both G proteins are linked to the same receptor. By the calculation above (with *n* = 2), we have

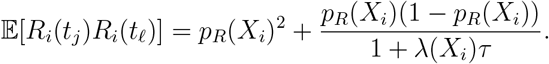

Therefore,

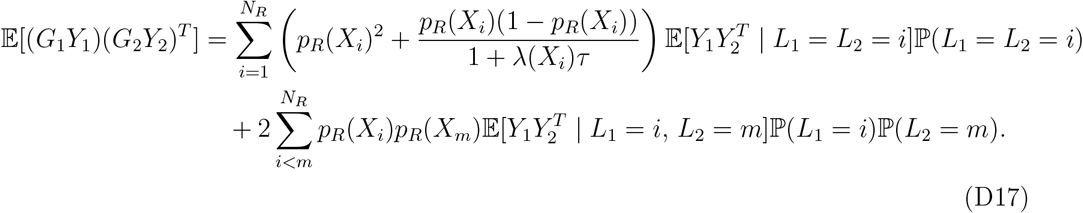

Putting this all together, we obtain

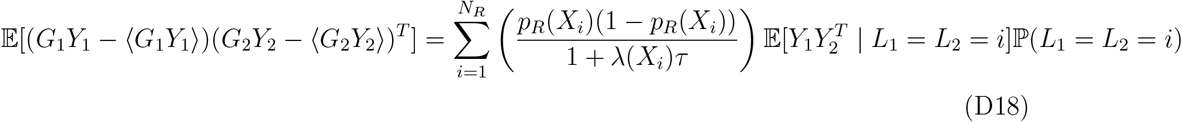

and the covariance matrix Σ_*G*_ = 𝔼 [(*V*_*G*_ − (*V*_*G*_))(*V*_*G*_ − (*V*_*G*_))^*T*^] is: Σ_*G*_ = *N*_*G*_

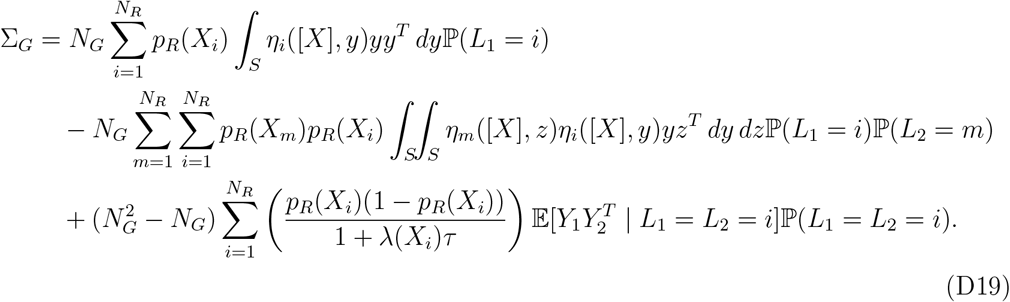

By definition of *η*_*i*_,

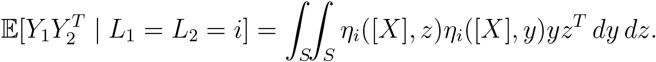

#### 5. Derivation of Proposition D.2

For any continuous function *f* on the sphere *S*, when *N*_*R*_ is large enough we have

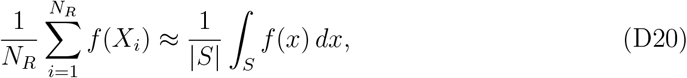

with high probability. This is justified by the law of large numbers, since the receptor positions {*X*_*i*_} are independent. Using this and the approximations (D7) and (D8), we have

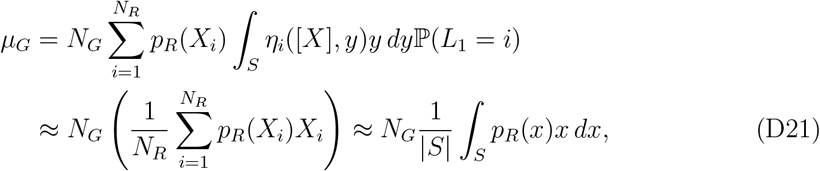

and similarly,

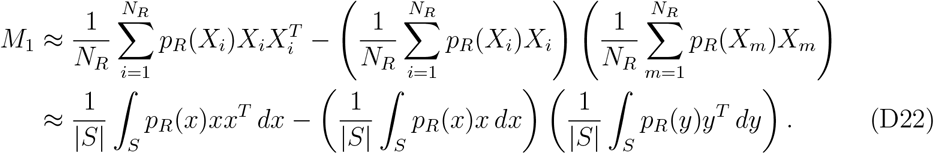

Since *L*_1_ and *L*_2_ are independent, ℙ (*L*_1_ = *L*_2_ = *i*) = ℙ (*L*_1_ = *i*) ℙ (*L*_2_ = *i*) ≈ (*N*_*R*_)^−2^. Therefore, using the same reasoning as for *M*_1_, we have

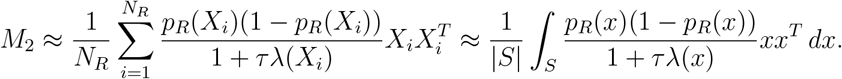

## Appendix E: Local concentration noise versus direction noise

Here is a simple illustration of the phenomenon that in the ratiometric model the gradient of active *G* protein may be stronger than in the classical model even while the local noise in active G protein may be greater for the ratiometric model. Thus, high-variability in local concentration of active G protein is not necessarily an indicator of poor direction estimation. This is because direction estimation involves a comparison across multiple different regions of the cell surface.

Imagine a greatly simplified, one-dimensional model where the cell has a front and a back: at the front there are *N*_*R*_ receptors and at the back there are *N*_*R*_ receptors. Let *E* ∈ (0, 1*/*2) be a small parameter representing a small gradient of ligand concentration (toward the front end). Let us suppose that 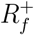 and 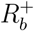 are independent random variables with distributions

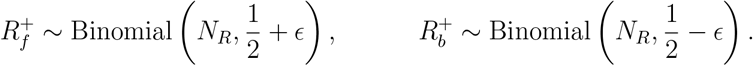

These 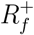 and 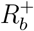 represent the number of active receptors at the front and back of the cell, respectively. This statistical model is equivalent to saying that the *N*_*R*_ proteins at the front end have independent states, each being active with probability 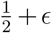. Similarly, at the back end, the probability of a receptor being active is 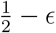. Then define

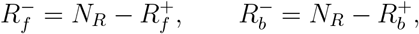

which represent the number of inactive receptor at the front and back of the cell, respectively. The mean receptor levels are

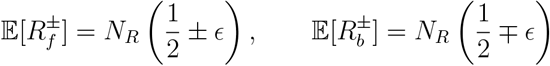

and the variances are

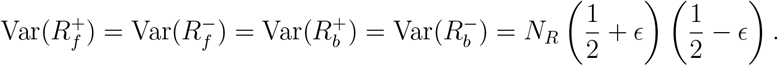

So, with *E* > 0, the receptor gradient has a small bias toward the front end:

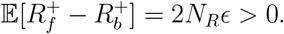

In addition to receptor, there are G proteins at the front and back. Suppose there are *N*_*G*_ proteins at the front and *N*_*G*_ proteins at the back. Random variables 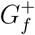 and 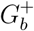 will be the number of active G proteins at the front and back, respectively. Given 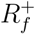 and 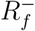, G proteins at the front end will be active with probability that depends on 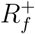 and 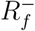; what distinguishes the ratiometric from classical models is how this probability depends on 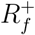 and 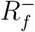. In the ratiometric model, the steady-state probability that a G protein at the front end is active is

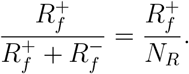

(Rate constants are normalized to one.) In the classical model, the inactivation rate is taken to be constant and uniform across the cell (same at the front and back). Therefore, in the classical model, the steady state probability that a G protein at the front end is active has the form

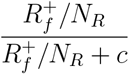

for some constant *c* > 0. Similar statements hold at the back end of the cell, with 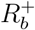 replacing 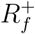. In either case, classical or ratiometic, the probability that a G protein is active, is a function of the local fraction of active receptor.

In view of this, we take 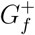 and 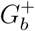 to be independent random variables having the following distribution: given 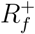 and 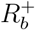,

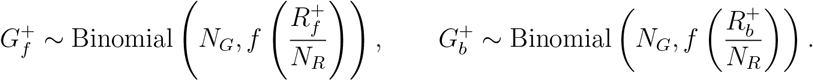

where

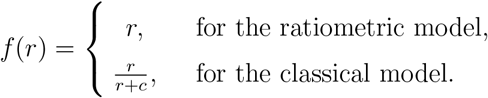

This statistical model is equivalent to saying that the *N*_*G*_ proteins at the front end have independent states, each being active with probability 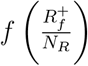. And similarly for the proteins at the back end.

To compare the two models, we choose the constant *c* so that the mean total activation rate of G protein is the same. The mean total activation is

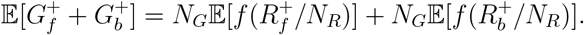

For the ratiometric model, the function *f* is linear (*f* (*r*) = *r*), so this can be computed explicitly:

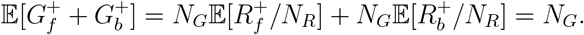

Therefore, the matching condition requires that the constant *c* (in the classical model) be chosen so that

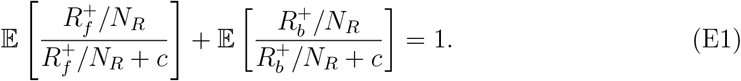

This relation for *c* cannot be solved explicitly. Nevertheless, we claim that when *N*_*R*_ is large and *E* is small, *c* ≈ 1*/*2. To see this, note that when *N*_*R*_ is large, the distributions of *R*_*f*_ */N*_*R*_ and *R*_*b*_*/N*_*R*_ are very concentrated around 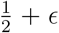 and 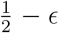, respectively. Therefore, when *ϵ* > 0 is small, (E1) implies that

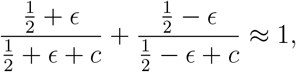

which implies that *c* ≈ 1*/*2 for small *ϵ*.

In the case of the classical model, *f* is nonlinear, and this makes it impossible to calculate explicitly the mean and variance of 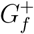 and 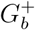 in that case. However, as we explained above, when *N*_*R*_ is large and *ϵ* is small, the constant *c* is approximately 1*/*2, and the function *f* may be approximated well by its linear expansion at the value *r* = 1*/*2:

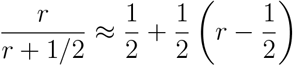

This motivates the following modification, which makes calculation explicit in both models: Given 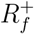 and 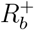, let

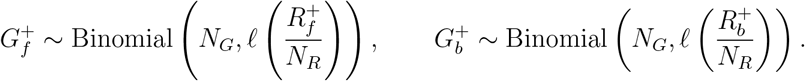

where, instead of *f* defined above, we use the function

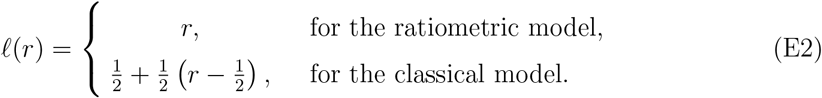

This *ℓ* is linear in both cases, and it has the form

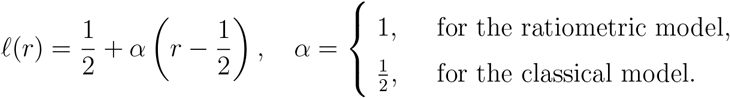

With this modification, the means and variances of 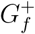 and 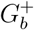 are computable in both models. We compute:

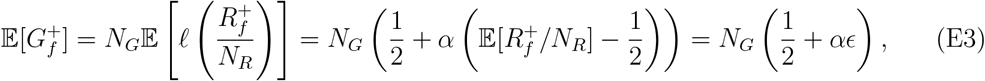

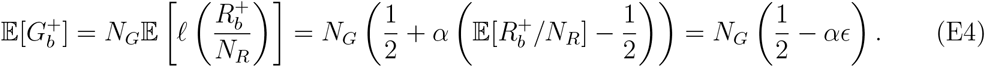

In particular, the total activation level is the same for all *α*:

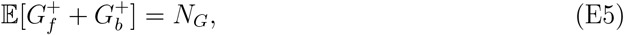

while the mean front-to-back gradient in active G protein is

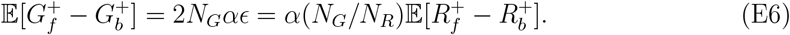

Recall that the ratiometric case corresponds exactly to *α* = 1, while the classical case corresponds to *α* = 1*/*2, so the mean gradient is twice as large in the ratiometric case.

Now, let us consider how the variances depend on *α*, especially the cases *α* = 1 (ratio-metric) and *α* = 1*/*2 (classical). The variance in active G protein, at both the front and back end, can be computed explicitly:

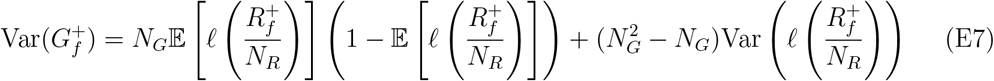

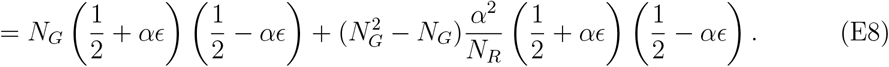

So, the coefficient of variation of 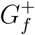 is

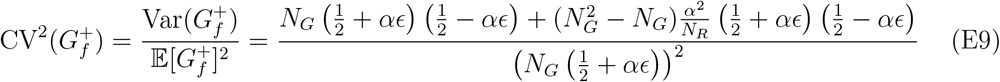

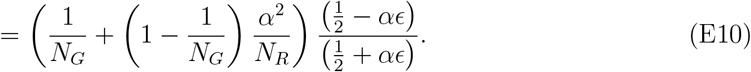

A similar compuation shows that at the back end we have

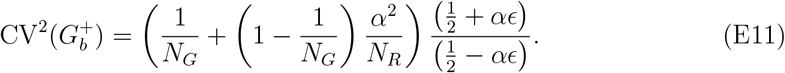

For small *ϵ*, this is approximately

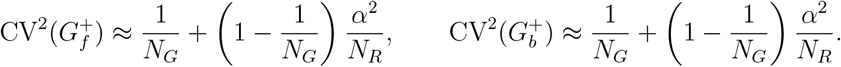

In particular, these quantities increase with *α* for *α* ∈ [0, 1]. On the other hand, the mean front-to-back gradient

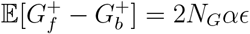

also increases with *α*. This shows that the directional signal gets stronger with larger *α*, even though the local noise levels (measured by coefficients of variation 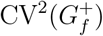 and 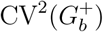) are larger with larger *α*. In particular, this illustrates the phenomenon that with the ratiometric model (*α* = 1), the local noise in 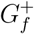 and 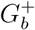 may be greater than in the classical model (*α* = 1*/*2), even while the front-to-back gradient in *G* protein is stronger for the ratiometric model.

This phenomenon can be seen more clearly from examining the joint distributions of 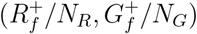 and 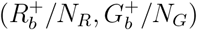 for the two models, see Figure 11. In Figure 12, we plot histrograms representing the marginal distributions of 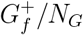 and 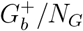 for the two models. The marginal distributions of 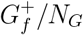 and 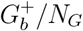 have higher variance for the ratiometric model, than for the classical model. Figure 13 shows a histrogram representing the distribution of active G protein gradient 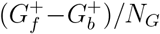 for both models. We observe that the distribution of this gradient is biased to the right (front end) for the ratiometric model, as explained in the analysis above. From Figure 11, we see how the different activation functions for the ratiometric vs. classical models leads to a bias in the G protein gradient from front to back, producing stronger (steeper) gradient in the ratiometric case, even while the variance in local G protein activation is higher with the ratiometric model.

**FIG. 11.**
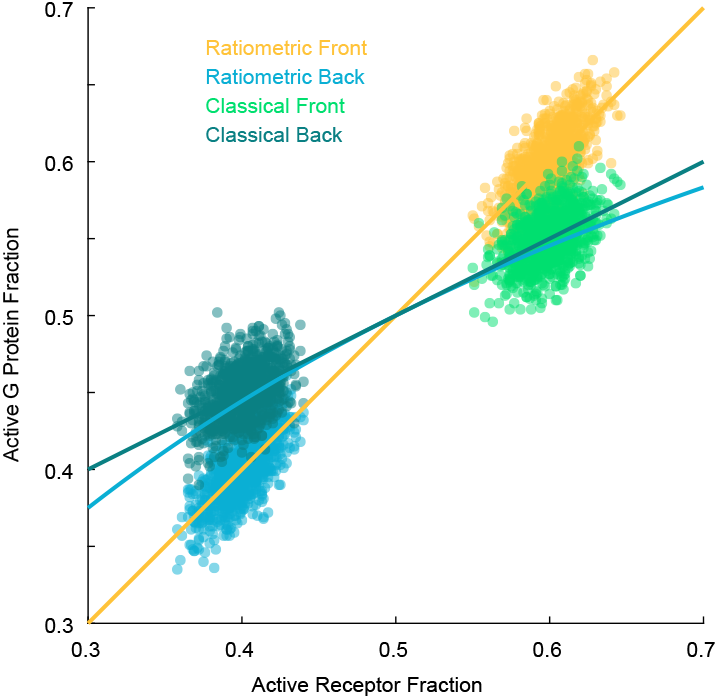
Empirical joint distribution of 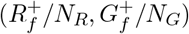 and 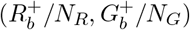, the active receptor and active G protein fractions at front and back of cell. Front of cell: ratiometric (yellow) and classical (green). Back of cell: ratiometric (blue) and classical (dark green). *N*_*R*_ = *N*_*G*_ = 1000. *ϵ* = 0.1. The red line is the (linear) activation function ℓ (*r*) = *r*, which applies to the ratiometric case. The blue and black lines are the activation functions *r/*(*r* + 1*/*2) and its linearization 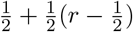, which applies to the classical model (see equation (E2)).

**FIG. 12.**
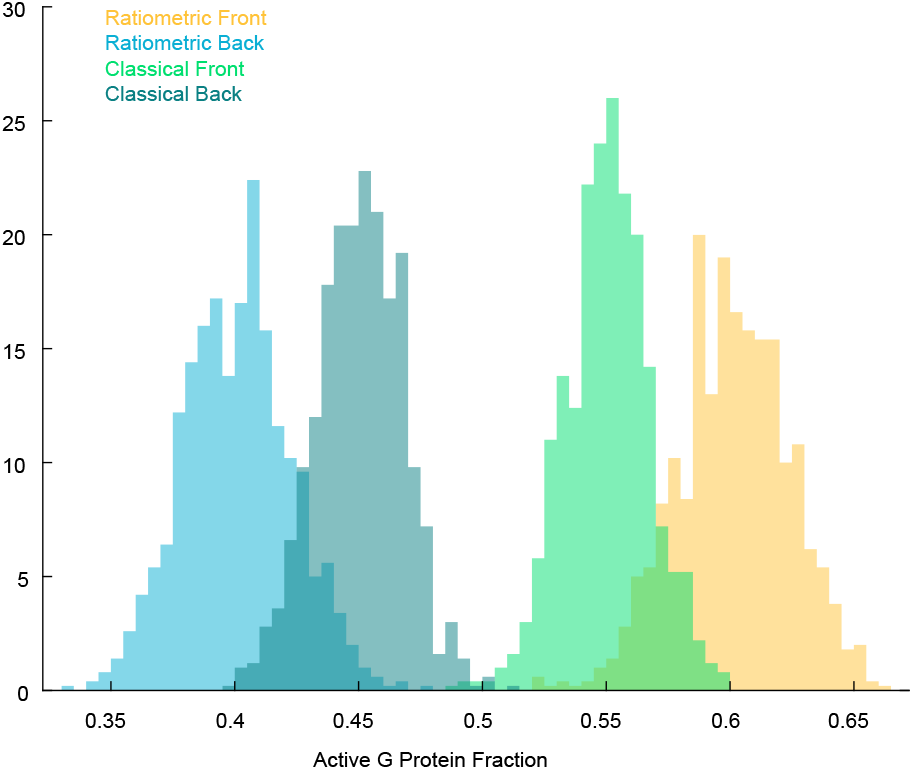
Distributions (histograms) of 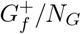 and 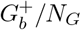, the active G protein fraction at the front and back of cell. Front of cell: ratiometric (yellow) and classical (green). Back of cell: ratiometric (blue) and classical (dark green). These correspond to marginal distributions from the joint distribution plot above, Figure 11. *N*_*R*_ = *N*_*G*_ = 1000. *E* = 0.1

**FIG. 13.**
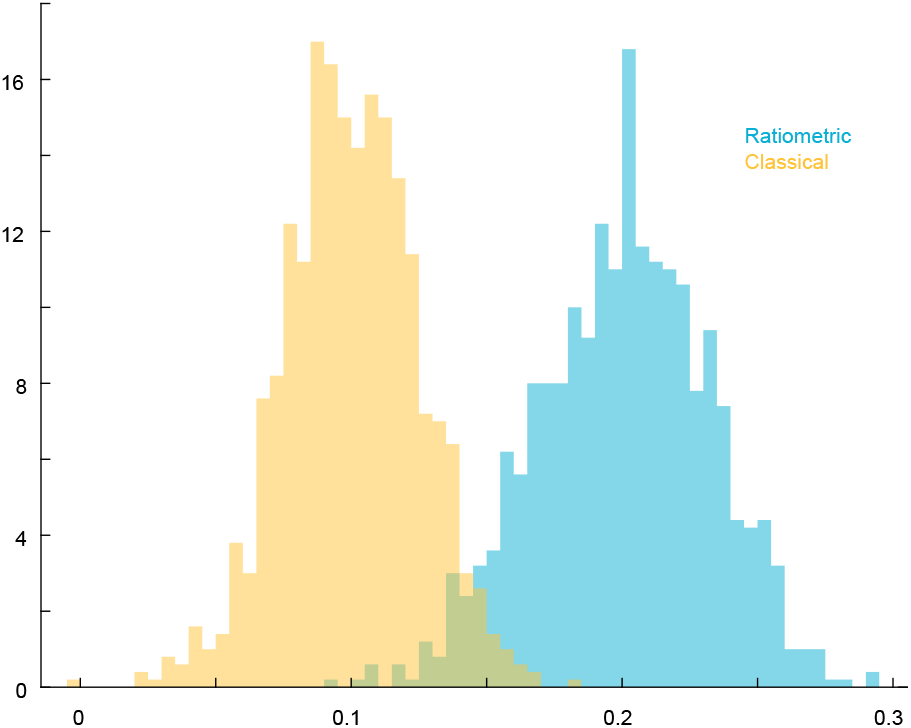
Distributions (histograms) of 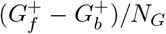, the active G protein gradient from front to back of cell, for Ratiometric (blue) and Classical (yellow).

## Appendix F: Classical model

Here we explain why the classical model (nonratiometric) performs more poorly than the ratiometric model. Recall that 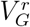 and 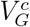 refer to the vector (B4) under the ratiometric and classical models, respectively. We have seen that the ratiometric model 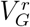 performs particularly well compared to *V*_*RL*_ and *V*_*C*_ when the time scales are such that the G proteins are approximately independent, and *N*_*G*_ > *N*_*R*_. In principle, the classical model estimate 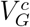 might also perform well in this scenario, yet our simulations show that this is not the case. An important point derived above is that for the ratiometric model, with *N*_*R*_ sufficiently large, the mean resultant vector is

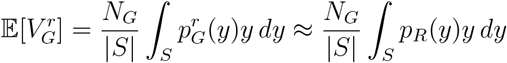

where 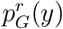 is the probability that a G protein at location *y* is active (under the ratiometric model); *p*_*R*_(*y*) is the probability that a receptor at *y* is active. In particular, the mean vector 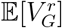 does not depend on λ_0_*τ*_dif_, where *τ*_dif_ = *τ* is the diffusive encounter time scale from (D1), and 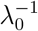 is the time scale of receptor switching. For the classical model, there is an additional time scale 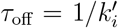, where 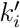 is the rate at which G proteins deactivate randomly in the classical model. For the classical model, the mean vector 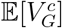 depends on all of these three time scales *τ*_off_, *τ*_dif_, and 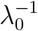 in a way that makes the classical model less effective at direction estimation than the ratiometric model.

For the classical model, let 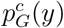 denote the probability that a G protein at position *y* is active. The probability 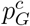 cannot be computed explicitly, but we analyze an approximate model that has similar features and gives insight into the way that 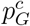 depends on the time scales. As we will show, the gradient of 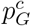may be much flatter than that of *p*_*R*_ or 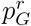. In the classical model, the state *G*(*t*) of a G protein at time *t* and position *Y* (*t*) = *y* is active if and only if that G protein encountered an active receptor (possibly multiple) at some time between time *t* − *T* and *t*, where the random time *T* has the Exponential distribution with mean *τ*_off_. Otherwise, that protein is inactive. The number of receptors that a G protein encounters before deactivation is random and it depends in a complicated way on the receptor configuration, but it is reasonable to suppose that random number *N* of distinct encounters has mean 𝔼 [*N*] = *τ*_off_*/τ*_dif_, where

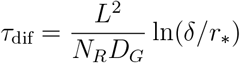

is the diffusive encounter time scale (see (D1)). Indeed, looking backward in time we may assume that the encounters between a G protein and receptors occur as a Poisson arrival process at rate 1*/τ*_dif_ (times between encounters are approximately exponentially distributed with mean *τ*_dif_). Thus, *N* has the distribution of *A*_*T*_, where *A*_*t*_ is such a Poisson arrival process and *T* ~ Exponential(1*/τ*_off_) is independent of the process *A*. This implies that the number *N* of distinct receptor encounters has the (shifted) Geometric distribution:

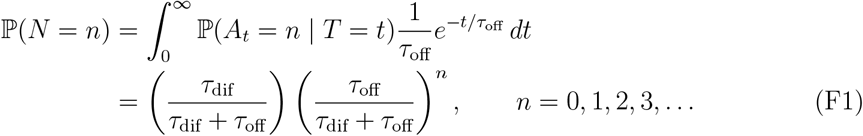

with mean 𝔼 [*N*] = *τ*_off_*/τ*_dif_. This distribution is in close agreement with our simulations; see Figure 14 where we show numerical computation of the empirical distribution of distinct receptors encountered by a G protein before its state resets (in the classical model).

**FIG. 14.**
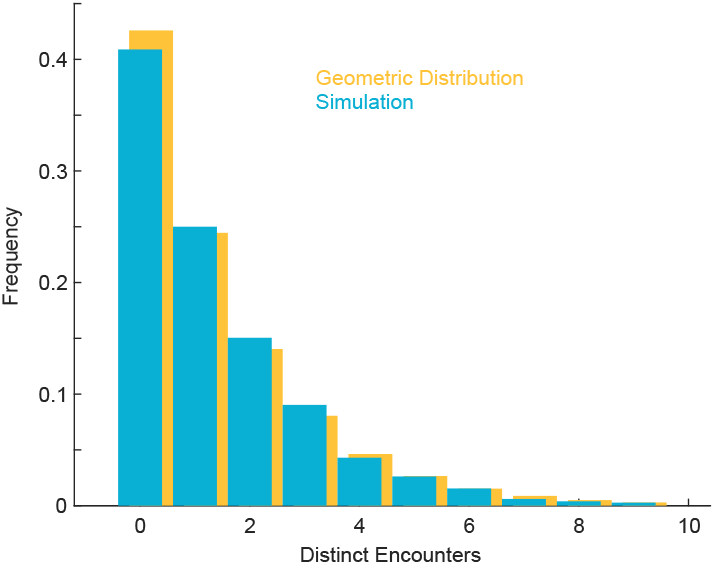
Blue: Empirical distribution of *N*, the number of distinct receptor encounters by a G protein since its most recent state-reset time. *N*_*R*_ = 375, *D*_*G*_ = 0.002(*µm*)^2^*/sec, k*_*go*_ = 0.0083(*sec*)^−1^. The mean number of encounters is 𝔼 [*N*] = 1.28. Fraction of proteins not encountering a receptor: 0.42. Mean time to first encounter: 79.9. 20,000 proteins were simulated, for a single realization of the receptor configuration. **Yellow:** For comparison, the Geometric distribution given by (F1).

An important consequence of this is that the probability that a G protein encounters zero receptors between resetting events is:

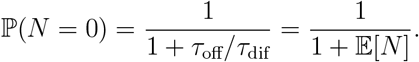

Such G proteins carry zero information about the receptor states, and thus zero information about the pheromone gradient or local concentration. This is an important difference between the classical and ratiometric models: in the classical model, at a given time *t*, some fraction (approximately 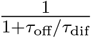) of proteins carry zero information about the local pheromone concentration because they have not encountered any receptor (active or inactive) since the most recent state resetting event; in the ratiometric model, however, every G protein carries some information about the local pheromone concentration, via the state of the most recently encountered receptor. Note that ℙ (*N* = 0) does not depend in any way on the pheromone concentration; 𝔼 [*N*] depends on the density of receptors, but not on their states.

The presence of “uninformative” G proteins (in the classical model) carrying no information about the pheromone gradient leads to what we call a flattening of the gradient of 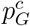 relative to *p*_*R*_. Assuming encounters are nearly instantaneous, the probability that a given encounter at location *x* activates the G protein is approximately *p*_*R*_(*x*). Assuming that the *N* encounters are local, then the probability that a G protein at *x* (under classical model) will be active is then

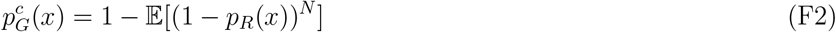

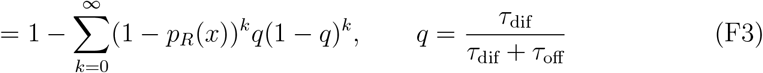

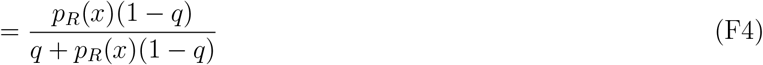

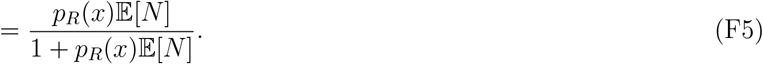

Thus the relationship between 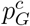 and *p*_*R*_ depends on the mean number of receptor encounters 𝔼 [*N*]. In Figure 15 we plot 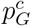versus *p*_*R*_ for different values of 𝔼 [*N*]. When *τ*_off_*/τ*_diff_ = 𝔼 [*N*] < < 1, most G proteins will be inactive, and (F5) shows that

**FIG. 15.**
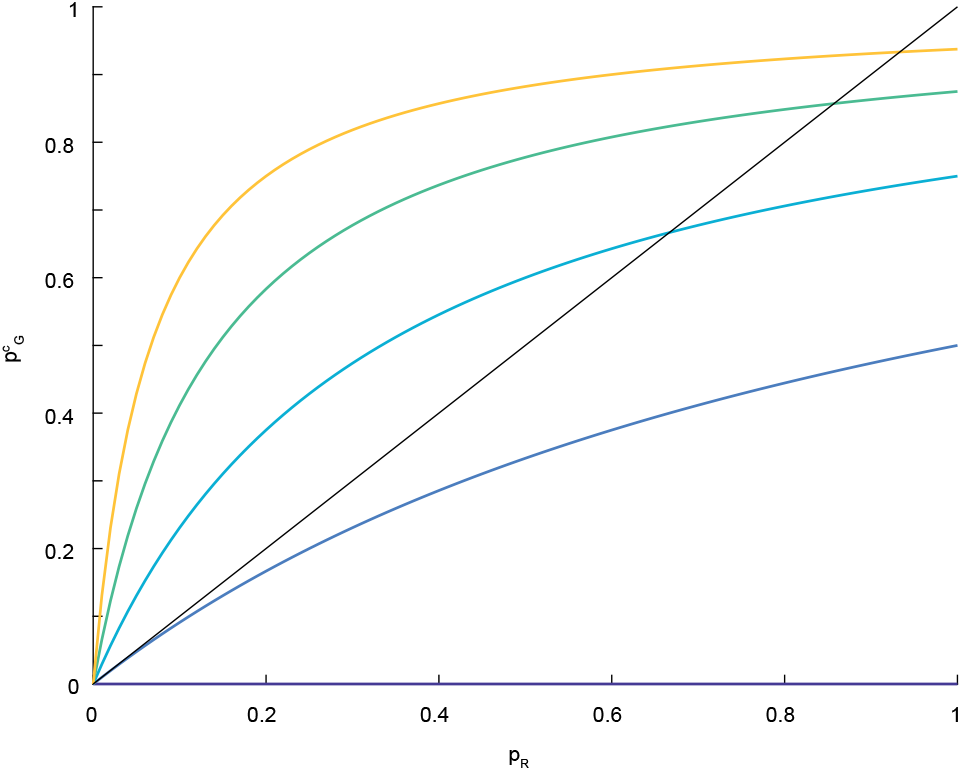
Graph of 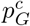 versus *p*_*r*_, for different values of 𝔼 [*N*] = 2^*k*^ − 1, *k* = 0, …, 4. For comparison, the straight line has slope 1.

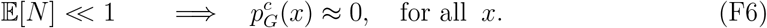

However, when *τ*_off_*/τ*_diff_ = 𝔼 [*N*] >> 1 and *p*_*R*_ not too small, there is saturation – most G proteins will be active, and (F5) shows that

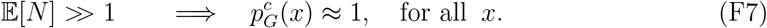

The gradient of 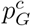 is

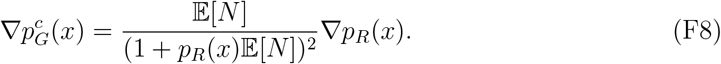

The prefactor in front of ∇*p*_*R*_ can be larger or smaller than 1. We have

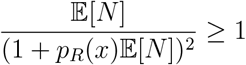

if only if 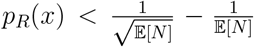. In particular, if 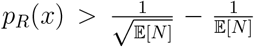, then there will necessarily be a flattening effect whereby the prefactor in (F8) is less than 1. Our earlier analysis shows that for shallow gradients, the NSR is smallest when *C*_0_ ≈ *K*_*d*_, which implies *p*_*R*_ is near 1*/*2. In this regime, 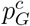 will be flatter than *p*_*R*_, as shown in Figure 15. In particular, 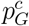 is flatter than 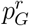, since for the ratiometric model 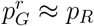 regardless of the time scales. The flattening of 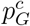 versus *p*_*R*_ and 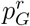 implies less accuracy (higher NSR) for the classical method, since the NSR scales inversely with the gradient squared (i.e. as 1*/b*^2^ in (B12).)

## Appendix G: Captions for Movies

**Movie S1**. Temporal dynamics of receptor and G-protein vector orientations during Classical gradient sensing. **Top left:** Full simulation showing all receptor and G-protein positions; receptors are color-coded by activation state. **Top middle:** Vector orientations for active (yellow), inactive (gray), and combined (blue) receptor populations. **Top right:** Vectors for the combined receptor (blue) and G-protein (green) populations. The black horizontal line denotes the true gradient direction. **Bottom:** Time traces showing the angle between the pheromone gradient and the receptor (green) and G-protein (cyan) vectors. NG = 2500, NR = 375, *D*_*m*_ = 0.002 *µm*^2^*/s, k*_*gi*_ = 0.0083 *s*^−1^, gradient = 8 to 4 nM. https://youtu.be/beVWFQ6Jzdg

**Movie S2**. Temporal dynamics of receptor and G-protein vector orientations during Classical gradient sensing. **Top left:** Full simulation showing all receptor and G-protein positions; receptors are color-coded by activation state. **Top middle:** Vector orientations for active (yellow), inactive (gray), and combined (blue) receptor populations. **Top right:** Vectors for the combined receptor (blue) and G-protein (green) populations. The black horizontal line denotes the true gradient direction. **Bottom:** Time traces showing the angle between the pheromone gradient and the receptor (green) and G-protein (cyan) vectors. NG = 2500, NR = 1500, *D*_*m*_ = 0.002 *µm*^2^*/s, k*_*gi*_ = 0.0357 *s*^−1^, gradient = 8 to 4 nM. https://youtu.be/nvEN7Sysycc

**Movie S3**. Temporal dynamics of receptor and G-protein vector orientations during Classical gradient sensing. **Top left:** Full simulation showing all receptor and G-protein positions; receptors are color-coded by activation state. **Top middle:** Vector orientations for active (yellow), inactive (gray), and combined (blue) receptor populations. **Top right:** Vectors for the combined receptor (blue) and G-protein (green) populations. The black horizontal line denotes the true gradient direction. **Bottom:** Time traces showing the angle between the pheromone gradient and the receptor (green) and G-protein (cyan) vectors. NG = 2500, NR = 6000, *D*_*m*_ = 0.002 *µm*^2^*/s, k*_*gi*_ = 0.1687 *s*^−1^, gradient = 8 to 4 nM. https://youtu.be/egZNPnUeXfM

**Movie S4**. Temporal dynamics of receptor and G-protein vector orientations during Ratiometric gradient sensing. **Top left:** Full simulation showing all receptor and G-protein positions; receptors are color-coded by activation state. **Top middle:** Vector orientations for active (yellow), inactive (gray), and combined (blue) receptor populations. **Top right:** Vectors for the combined receptor (blue) and G-protein (green) populations. The black horizontal line denotes the true gradient direction. **Bottom:** Time traces showing the angle between the pheromone gradient and the receptor (green) and G-protein (cyan) vectors. NG = 2500, NR = 375, *D*_*m*_ = 0.002 *µm*^2^*/s*, gradient = 8 to 4 nM. https://youtu.be/B3Dg8SMIhEE

**Movie S5**. Temporal dynamics of receptor and G-protein vector orientations during Ratiometric gradient sensing. **Top left:** Full simulation showing all receptor and G-protein positions; receptors are color-coded by activation state. **Top middle:** Vector orientations for active (yellow), inactive (gray), and combined (blue) receptor populations. **Top right:** Vectors for the combined receptor (blue) and G-protein (green) populations. The black horizontal line denotes the true gradient direction. **Bottom:** Time traces showing the angle between the pheromone gradient and the receptor (green) and G-protein (cyan) vectors. NG = 2500, NR = 1500, *D*_*m*_ = 0.002 *µm*^2^*/s*, gradient = 8 to 4 nM. https://youtu.be/9SNJQRN9yAQ

**Movie S6**. Temporal dynamics of receptor and G-protein vector orientations during Ratiometric gradient sensing. **Top left:** Full simulation showing all receptor and G-protein positions; receptors are color-coded by activation state. **Top middle:** Vector orientations for active (yellow), inactive (gray), and combined (blue) receptor populations. **Top right:** Vectors for the combined receptor (blue) and G-protein (green) populations. The black horizontal line denotes the true gradient direction. **Bottom:** Time traces showing the angle between the pheromone gradient and the receptor (green) and G-protein (cyan) vectors. NG = 2500, NR = 6000, *D*_*m*_ = 0.002 *µm*^2^*/s*, gradient = 8 to 4 nM. https://youtu.be/JxBwMO2otD8

